# The K^+^ exchange antiporter 3 senses the chloroplast energy status to synchronize photosynthesis

**DOI:** 10.1101/2022.12.19.521033

**Authors:** Michal Uflewski, Tobias Rindfleisch, Kübra Korkmaz, Enrico Tietz, Sarah Mielke, Viviana Correa Galvis, Beatrix Dünschede, Marcin Luzarowski, Aleksandra Skirycz, Markus Schwarzländer, Deserah D. Strand, Alexander P. Hertle, Danja Schünemann, Dirk Walther, Anja Thalhammer, Martin Wolff, Ute Armbruster

## Abstract

Plant photosynthesis contains two functional modules, the light-driven reactions in the thylakoid membrane and the carbon-fixing reactions in the chloroplast stroma. In nature, light availability for photosynthesis often undergoes massive and rapid fluctuations. Efficient and productive use of such variable light supply requires an instant crosstalk and rapid synchronization of both functional modules. Here, we show that this communication involves the stromal exposed regulatory C-terminus (RCT) of the thylakoid K^+^-exchange antiporter KEA3. RCT-mediated control of KEA3 contributes to the balance between light capture and photoprotection. By combining *in silico, in vitro*, and *in vivo* approaches, we demonstrate that the RCT senses the energy state of the chloroplast in form of both, phosphorylation and redox potential, in a pH-dependent manner and regulates KEA3 activity in response. Together our data pinpoint a regulatory feedback loop by which the stromal energy state orchestrates light capture and photoprotection via KEA3.

## Introduction

In dense plant habitats, such as crop fields, sunlight availability for photosynthesis can undergo massive changes on short time scales^1,2^. In such dynamic light environments, the photoprotective energy-dependent quenching (qE) is crucial for avoiding light stress^3^. However, its rather slow relaxation kinetics limit crop photosynthesis and yield ^4,5^. Thus, manipulating qE dynamics holds great promise for enhancing crop photosynthesis in the field.

The thylakoid K^+^ exchange antiporter 3 (KEA3) is a key regulator of qE relaxation^6-11^. By exporting protons from the lumen in exchange for stromal K^+^, KEA3 decreases lumen pH-dependent qE. Under excess light intensities, when high qE is needed for photoprotection, KEA3 is inactivated. This light intensity-dependent KEA3 inactivation was shown to involve a C-terminal soluble domain^7,9^. Lack of this regulatory C-terminus (RCT) led to increased damage of both photosystems during a light stress treatment^9^. However, during the initial transfer from dark to high light, such RCT-less plants showed increased rates of photosynthesis. This finding introduced KEA3 as a promising candidate for manipulating qE dynamics to enhance photosynthesis and advocated for a detailed mechanistic understanding of RCT-mediated regulation^9^.

The RCT contains the conserved K^+^ transport nucleotide binding (KTN) domain, which in other transport systems was shown to gate K^+^-transport processes in response to nucleotide binding^12-16^. The light reactions of photosynthesis convert light energy into redox and phosphorylation potential by using the nucleotide couples NADPH, NADP^+^ and ATP, ADP, respectively^9,10^. Thus, ratios of these nucleotides reflect on the energy state of the chloroplast stroma and may serve as signals for the light-intensity dependent regulation of KEA3 activity. Another signature of photosynthetic activity, and therefore a potential signal, is the acidification of the lumen and simultaneous alkalization of the chloroplast stroma.

In the current work we combine *in silico, in vitro* and *in vivo* approaches to (i) show the RCT is exposed to the chloroplast stroma, (ii) characterize the response of stromal pH to changes in light intensity *in planta* by using a genetically encoded pH sensor, (iii) resolve the interaction of the RCT with nucleotides and effects of pH, and (iv) characterize the role of RCT nucleotide binding in qE dynamics.

## Results

### The regulatory KEA3 C-terminus is exposed to the chloroplast stroma

Previously, protease treatments of intact thylakoid membranes to resolve the localization of the KEA3 C-terminus had yielded inconclusive results^11,17^. To revisit the membrane topology of KEA and the localization of the KEA3 C-terminus, the self-assembling green fluorescent protein (saGFP) system was used, which yields fluorescence when both GFP fragments (saGFP1 and saGFP2) are co-localized in the same cellular compartment and spontaneously assemble^18^ (Fig. 1a). A protein fusion of KEA3 with the small GFP fragment (saGFP2) attached to the KEA3 C-terminus yielded fluorescence when co-expressed together with saGFP1 targeted to the stroma. This strongly suggested that the KEA3 C-terminus is localized at the stromal side of the thylakoid membrane (Fig. 1b, Supplemental Fig. 1). The co-expression of the same construct together with a cytosolic saGFP1 as well as a fusion protein of saGFP2 attached to the KEA3 N-terminus together with stromal saGFP1 instead yielded no clear GFP-fluorescence. A stromal localization of the regulatory C-terminus (RCT) was further supported by: (i) proteolytic digestion of stromal exposed proteins from intact thylakoids using additional lines (Supplemental Fig. 2) and (ii) association of stroma-targeted RCT with the thylakoid membrane as a function of KEA3 (Supplemental Fig. 3).

**Fig. 1.**
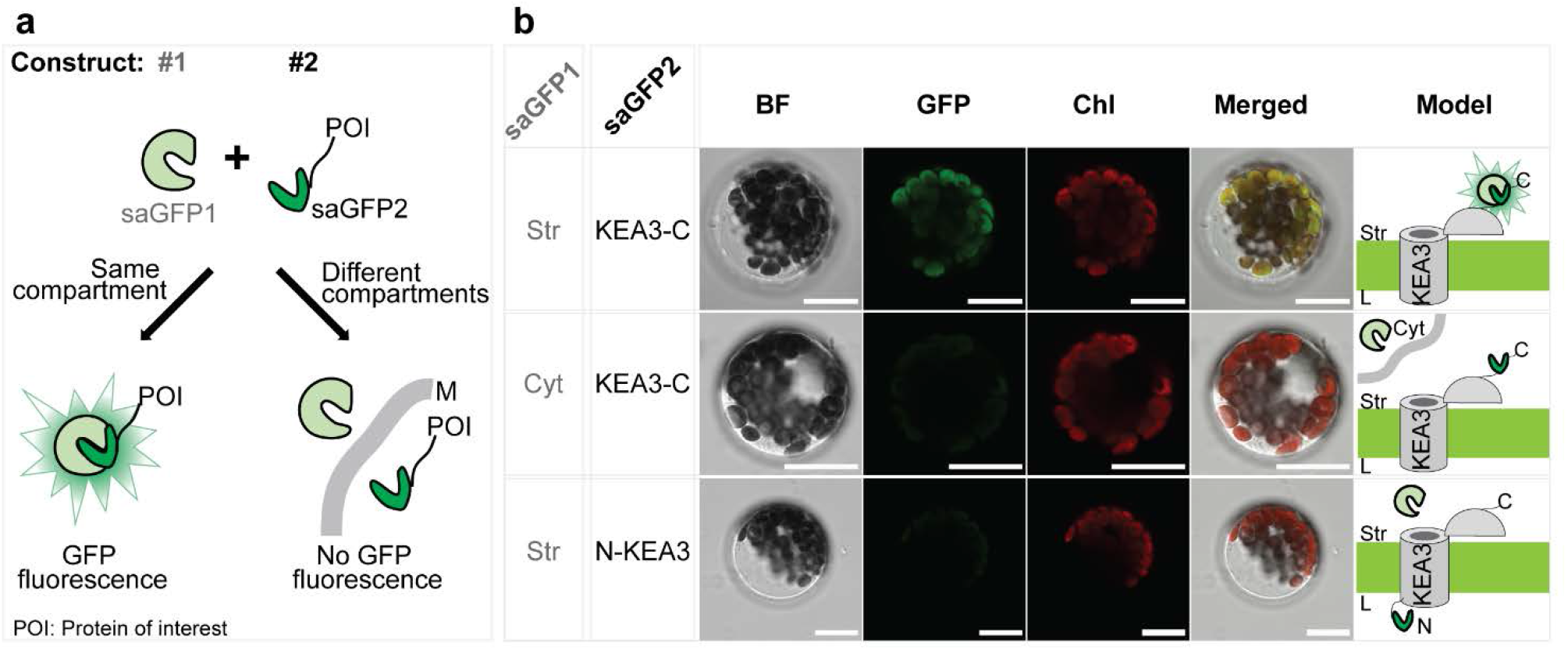
The regulatory KEA3 C-terminus is exposed to the chloroplast stroma. **a**, The self-assembling (sa) GFP system was used to determine the localization of the regulatory KEA3 C-terminus. The large saGFP1 fragment is targeted to a certain cellular compartment. The protein of interest (POI), in our case KEA3, is fused at the N- or C-terminus to the smaller saGFP2 fragment. If both saGFP fragments localize in the same cellular compartments they spontaneously assemble and yield GFP fluorescence (M, membrane). **b**, Transient co-expression of stromal targeted saGFP1 and a C-terminal saGFP2 fusion to KEA3 (KEA3-C) in Arabidopsis protoplasts results in GFP fluorescence. Cytoplasmic (Cyt) saGFP1 and KEA3-C as well as stromal saGFP1 and a N-terminal saGFP2 fusion to KEA3 (N-KEA3) do not yield GFP fluorescence. Scale bar, 15 µm; BF, bright field; Chl, chlorophyll fluorescence.

### Light intensity transitions induce marked changes in stromal pH

To elucidate stromal pH changes in response to light fluctuations and correlate them with changes in KEA3 activity, we employed plants expressing stroma-localized circular permuted yellow fluorescent protein (cpYFP) ^19^. By using a confocal microscopy set-up, we exposed dark acclimated leaf sections of cpYFP plants to fluctuations in light intensity. The analysis revealed that stromal pH responds to changes in light intensity in a highly dynamic fashion with a transient maximum in response to higher light intensities and a transient minimum in response to lower light intensities (Fig. 2a, Supplemental Fig. 4a). Steady state levels of stromal pH were reached ∼ 2 minutes after each light shift and differed significantly between the two light intensities. No such differences in fluorescence at the same detected wavelength window were observed in plants that carried the pH insensitive enhanced GFP (eGFP) in the stroma.

**Fig. 2.**
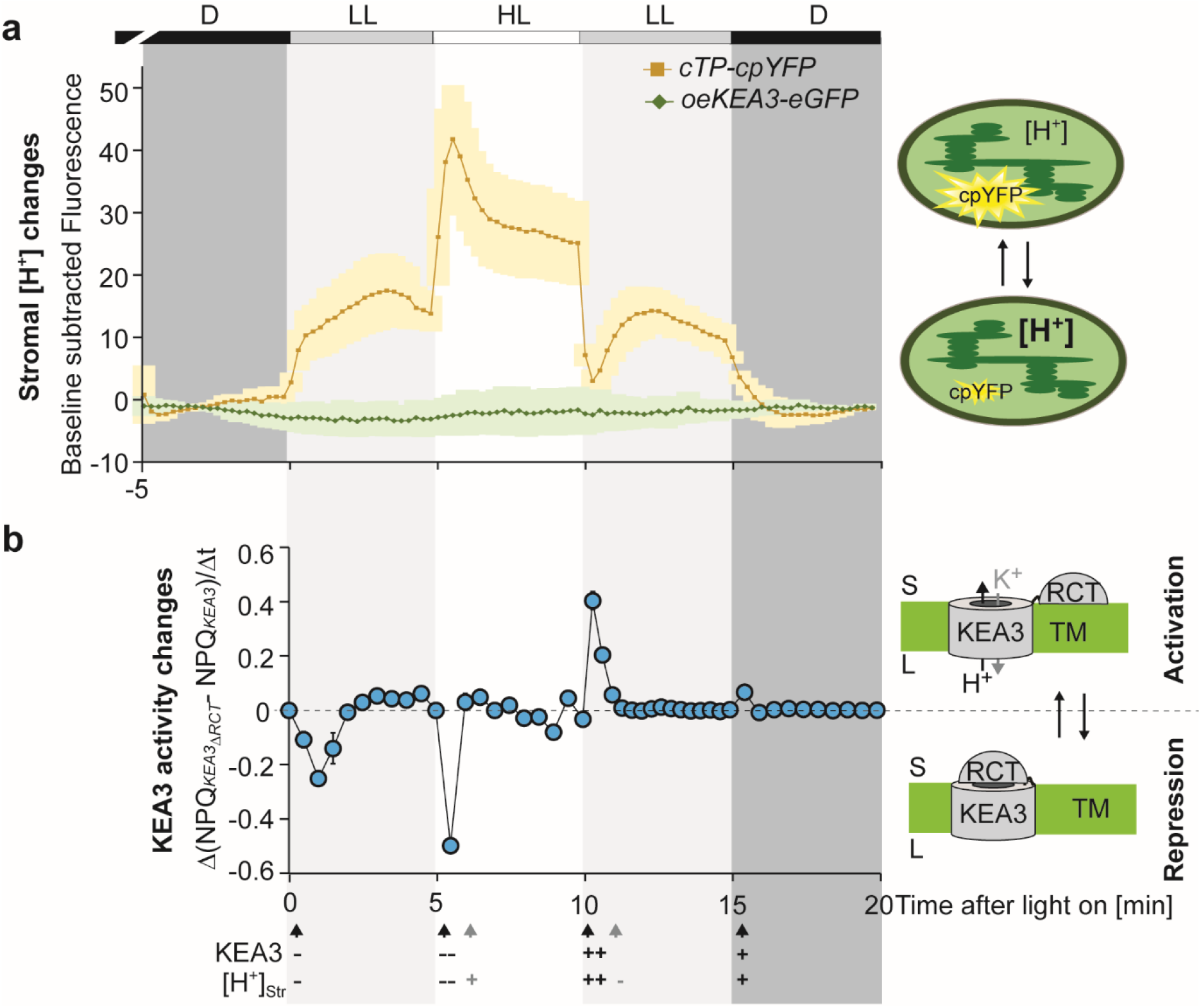
Stromal pH strongly responds to light intensity transients and transitions coincide with KEA3 regulation. **a**, Stromal pH changes in response to light intensity transitions were measured using plants expressing a stromal targeted circular permuted yellow fluorescent protein (cpYFP)^19^ and as control plants overexpressing a full-length KEA3 protein with C-terminal eGFP (oe*KEA3-eGFP*)^*17*^. Leaf sections were dark-acclimated (D, darkness) and examined by confocal microscopy using a light box, which supplied red light between 620 and 645 nm wavelengths at two different light intensities (low light, LL, 90, and high light, HL, 900 µmol photons m^-2^ s^-1^) during indicated time intervals. Fluorescence values shown were collected at 498-548 nm and subtracted from a dark baseline through the first and last minute of the measurements. Average is shown for n = 3 ± SD (Supplemental Fig. 4a for single traces and representative images). **b**, The regulatory C-terminus (RCT)-dependent difference in NPQ between two measuring time points is used as an approximation for RCT-dependent repression or stimulation of KEA3-mediated proton export from the lumen. NPQ during the same fluctuating light treatment as in (a) was determined of plants expressing *KEA3* (*KEA3-eGFP* in *kea3-1*) or *KEA3*_*ΔRCT*_ (*KEA3*_*ΔRCT*_*-eGFP* in *kea3-1*) from the KEA3 promoter9 and the difference between ΔNPQ (*KEA3*_*ΔRCT*_ -*KEA3*) between two measuring time points was calculated. Average is shown for n = 5 ± SD. Apparent simultaneous changes in KEA3 activity and stromal proton concentration are indicated by black arrows, stromal changes in proton concentration alone by grey arrows. The minus and plus signs indicate decreases and increases, respectively, with number of signs depicting the magnitude of change.

### RCT mediated KEA3 activity changes coincide with light induced stromal pH transitions

Next, we approximated RCT-dependent changes in KEA3 activity during light fluctuations from NPQ differences between plants with WT-like levels of either KEA3-eGFP or KEA3_ΔRCT_-eGFP^9^ (Supplemental Fig. 4b). An increase or decrease in the NPQ differences between two time points was interpreted as an RCT-dependent inactivation or activation of KEA3 activity, respectively (Fig. 2b). This analysis showed that RCT-dependent regulation of KEA3 correlates with rapid changes in stromal pH induced by light intensity transitions. Because not all changes in stromal pH were accompanied by changes in KEA3 activation, stromal pH alone, however, is unlikely to account for the full regulation of KEA3 activity via the RCT.

### KEA3 binds to ATP-linked beads

By using ATP and AMP-coupled agarose, we pulled down binding proteins from solubilized thylakoid membranes and tested for presence of KEA3 (Fig. 3a). KEA3 was detected on three of four different types of ATP-linked agarose beads but on none of the AMP-linked beads (Fig. 3b). We evaluated whether RCT mediates the binding of KEA3 to the ATP beads by using thylakoids of KEA3-eGFP and RCT-less KEA3_ΔRCT_-eGFP (expressed from the KEA3 promoter, Fig. 3c). The results showed that more KEA3 was pulled down from KEA3-eGFP harboring thylakoids, supporting that the RCT is involved in ATP-binding. Residual binding of KEA3_ΔRCT_-eGFP to the ATP-beads may be explained by additional binding sites located in the transport domain or the remaining 19 AA of the RCT that are still present in the KEA3_ΔRCT_-eGFP version retaining some affinity for ATP (Supplemental Fig. 5a). KEA3 pull-down by ATP-linked beads was observed at the two physiological pH values 6.8 and 8.0, which were used to approximate minimum and maximum pH of the stroma.

**Fig. 3.**
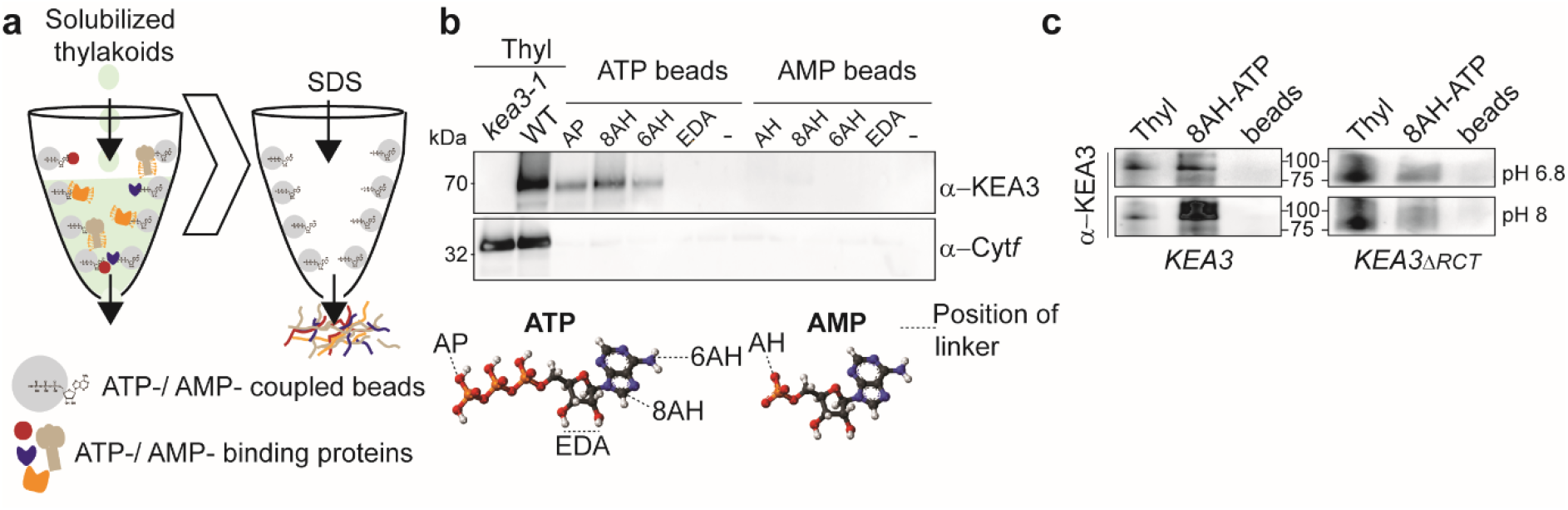
KEA3 is purified from solubilized thylakoid membranes by ATP-linked agarose beads. **a**, Scheme of experimental set-up. Thylakoid membranes were solubilized with ß-DM and incubated with ATP-/AMP-coupled agarose beads. After thoroughly washing off unbound proteins, ATP and AMP-binding proteins were eluted with SDS. **b**, Immunoblot analysis of *kea3-1* and WT thylakoids (WT) and the affinity pull down of four different kinds of ATP and AMP-linked beads from solubilized WT thylakoid membranes. The positions of the different linkers of the beads with the nucleotides are indicated below the immunoblot results. Specific KEA3 and Cyt*f* antibodies were used for hybridization. Cyt*f* shows specificity of the KEA3-ATP interaction, as different to KEA3 it does not bind to any of the different agarose beads. **c**, Affinity purification was performed on solubilized thylakoids of *KEA3* (*KEA3-GFP* in *kea3-1*) and *KEA3*_*ΔRCT*_ (*KEA3*_*ΔRCT*_*-GFP* in *kea3-1*) at pH 6.8 and 8. Immunoblots were hybridized with the specific KEA3 antibody.

### Recombinant RCT is monomeric

Structure prediction revealed that the RCT is composed of two Rossmann folds (Supplemental Fig. 5a). The KTN domain forming Rossmann fold 1 and the Rossmann fold 2 (RF2) are connected by an additional helix (connecting helix, CH). For further characterization of the RCT we performed size exclusion chromatography (SEC, Fig. 4a), static light scattering (SLS) for absolute molecular weight quantification and dynamic light scattering (DLS, Fig. 4b). Both SEC and DSL yielded a hydrodynamic radius of ∼ 3.8 nm for the RCT. The SLS measurement revealed the RCT to have a molecular mass of ∼30 kDa, which is close to the predicted mass of 32.3 kDa of an RCT monomer. The rather large hydrodynamic radius can be explained by the RCT not having a compact globular conformation. Far-UV CD experiments confirmed the RCT to contain large stretches of unstructured regions adding up to ∼40% of the entire protein (Supplemental Fig. 5b-c). Addition of an excess amount of ATP to the RCT did not change its monomeric status, but slightly decreased the hydrodynamic radius (Fig. 4b). To identify the ATP binding site of the RCT, we performed *in silico* docking (Supplemental Fig. 5d). This analysis indicated ATP binds in the vicinity of the first glycine (G65) of the glycine-rich region within the conserved KTN nucleotide binding loop of the Rossmann fold (Fig. 4c).

**Fig. 4.**
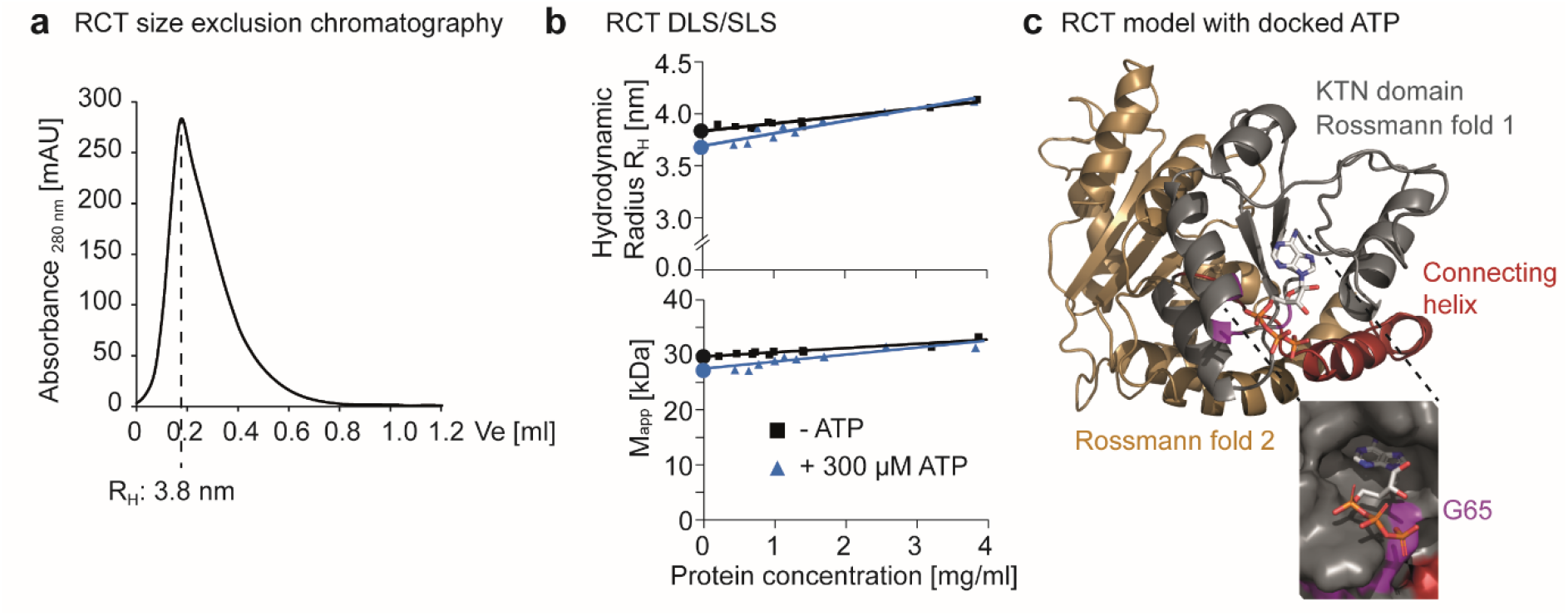
The KEA3 RCT binds ATP as a monomer. **a**, Size exclusion chromatography (SEC) profile of recombinant RCT after purification. Displayed is the elution volume (Ve) starting from the solvent front as determined by the peak volume of blue dextran elution. The SEC column was calibrated with globular protein standards and the hydrodynamic radius (R_H_) were calculated based on these standards. **b**, Dynamic light scattering (DSL, upper panel) and static light scattering (SLS, lower panel) were performed for total quantification of R_H_ and the apparent molecular mass (M_app_) of the RCT with and without 300 µM ATP by measuring different protein concentrations (indicated by symbols) and performing linear regression to extrapolate the intercept (indicated by circles). This analysis revealed the RCT being monomeric in solution with a larger hydrodynamic radius than expected from a globular protein. **c**, Predicted model of the RCT with docked ATP. The model predicts two Rossmann folds, 1 (KTN domain) and 2, which are colored in silver and gold, respectively. A helix connecting both Rossmann folds is depicted in red. The three glycines of the conserved GXGXXG glycine stretch are colored purple. The ATP is docked close to the first glycine, which in the recombinant RCT is located at amino acid residue position 65 (G65).

### MD simulations predict ATP binding to change RCT conformation

To further investigate the role of G65 in binding of ATP, we performed molecular dynamics (MD) simulations of wild-type RCT (RCTWT) and an RCT version, in which G65 was replaced by alanine (RCT_G65A,_ Fig. 5a). MD simulations of RCT_WT_ together with ATP confirmed a high frequency of ATP occupancy at a stretch of 12 AA including G65, while ATP presence at this site was largely interrupted in trajectories of the RCT_G65A_ mutant (Fig. 5b). We next had a look at the radius of gyration calculated from the model after MD simulation as a measure for global changes in RCT protein. The analysis indicated that ATP binding decreases the RCT_WT_ radius of gyration (Fig. 5c). The radii distributions calculated from the models after MD simulation were similar between RCT_G65A_ and RCT_WT_ + ATP with a pronounced peak at ∼22 Å. In the absence of ATP, the RCT_WT_ radii distribution showed a prominent maximum at ∼24 Å instead. To obtain a deeper understanding of the underlying structural changes that cause the simulated differences in dimensions, we compared secondary structures of the different models. By plotting the differences in secondary structure content to ligand-free RCT_WT_, we found distinct differences, which again were similar between RCT_WT_ + ATP and both RCT_G65A_ simulations. All three simulated models showed a loss of ß-sheet content at either side of and an increase in α-helical structures within the G65-containing ATP binding region as compared to the ligand-free RCT_WT_ (Fig. 5d). Additionally, they showed a strong increase in α-helical conformation at positions RCT166-178 and RCT216-227 within the second Rossmann fold.

**Fig. 5.**
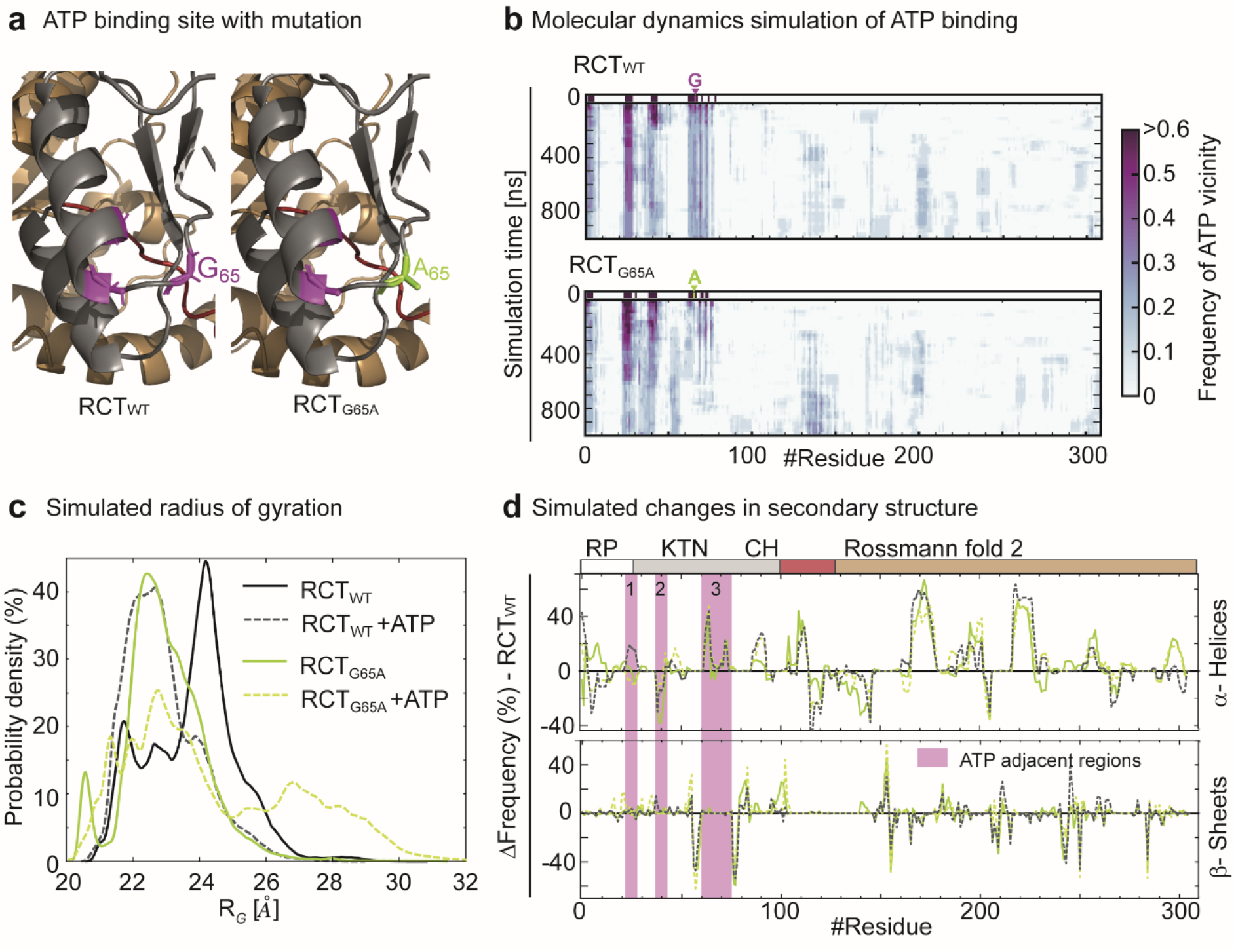
An RCT with single amino acid exchange loses ATP during simulation while mimicking an ATP-bound conformation. **a**, Model of the ATP binding site of wild-type RCT (RCT_WT_, left) and a variant carrying a G65A mutation (RCT_G65A_, right). The KTN domain is colored silver, the connecting helix (CH) red, the Rossmann fold 2 in gold and the glycines of the glycine-rich stretch in purple. **b**, Molecular dynamics (MD) simulations were performed on RCT_WT_ and RCT_G65A_ with ATP docked to its predicted binding site. The distance from the ATP molecule was calculated for each amino acid residue during the time course of the simulation. ATP spatial vicinity was defined by a distance of less than 7 Å from the ATP molecule. In the RCTWT simulation, a region containing the glycine stretch and thus G65 (highlighted in purple), remained in close contact with the ATP during simulation. The frequency of ATP vicinity decreased for the same region containing the single amino acid exchange of RCT_G65A_ over time leading to a loss of ATP vicinity for this region at the end of the simulation. Two further ATP adjacent regions were identified by the docking, which also lost ATP during simulation particularly in RCT_G65A_. Shown is relative ATP vicinity as calculated from n = 10 simulations. **c**, The probability density was calculated for radii of gyration from the MD simulation. The analysis shows that ATP binding to the RCT_WT_ leads to a decrease in protein radius, which is mirrored by the RCT_G65A_ mutation. **d**, Frequencies of α-helices and ß-sheets along the RCT protein sequence during MD simulations of RCT_WT_ and RCT_G65A_ with and without ATP are displayed as the difference to ligand-free RCT_WT_. The ATP adjacent regions 1-3 are shown in purple. Frequencies are calculated from n = 10 simulations. RP, recombinant part of the RCT. Same color code for simulations as shown in c.

### Binding of ATP and ADP involves the conserved glycine stretch of the C-terminal KTN domain

Next, recombinant RCT_G65A_ was produced alongside RCT_WT_ to experimentally characterize ligand interactions (Supplemental Fig. 6). Microscale thermophoresis (MST) analysis supported binding of ATP by RCT_WT_. It revealed ATP-induced fluorescence changes during MST were dependent on pH. Fluorescence differences were higher at pH 7.0 than at pH 8.0 (Fig. 6a). In line with the simulation results, RCT_G65A_ did not show any response to increasing concentrations of ATP during MST analysis, supporting that this RCT variant does not bind ATP. Because most of the MST traces did not follow a sigmoidal binding curve, but rather suggested two phases of rising fluorescence, we refrained from fitting the data to obtain a single dissociation constant.

**Fig. 6.**
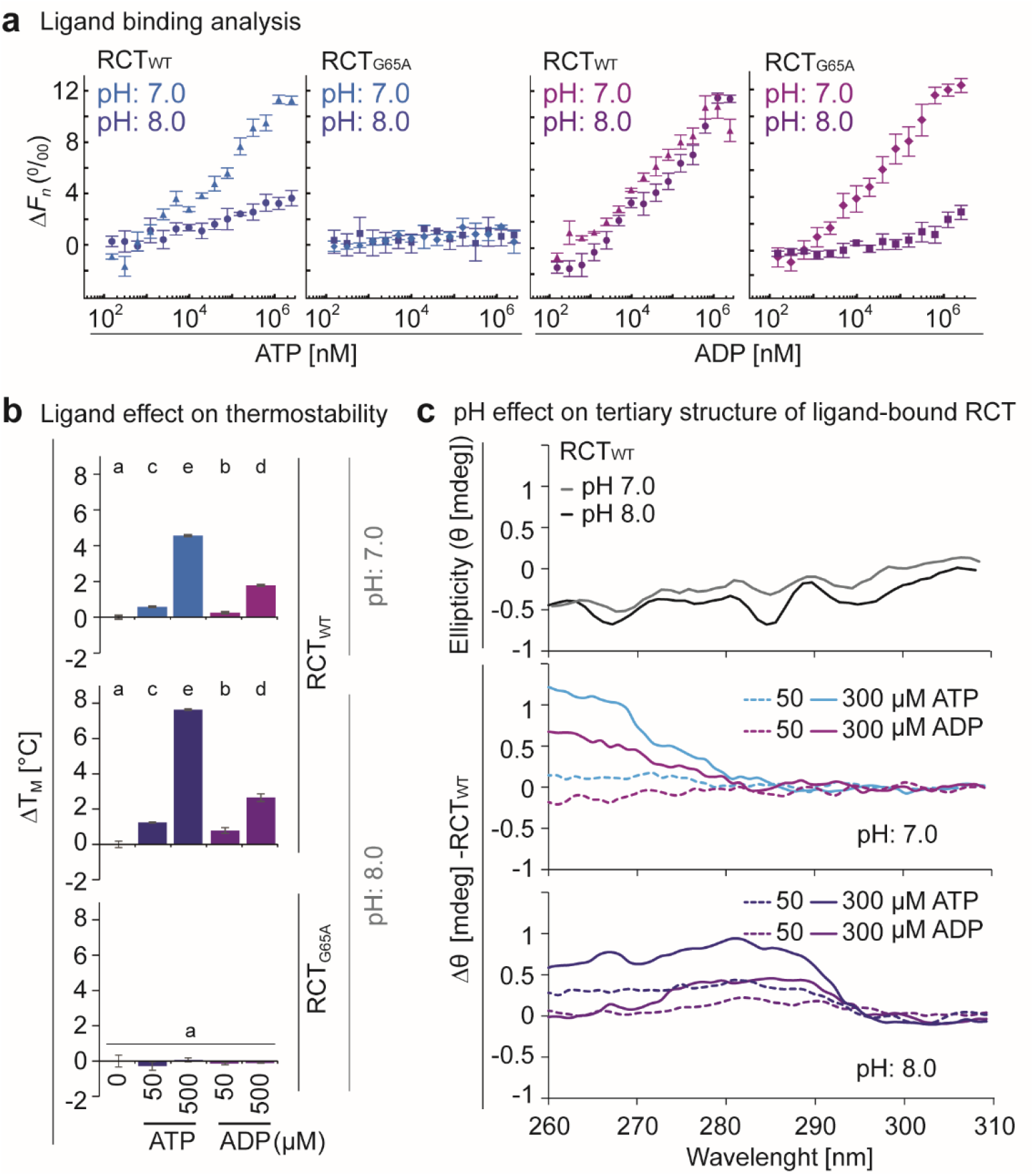
ATP and ADP binding to the KEA3 regulatory C-terminus induces pH-dependent conformational changes. **a**, The difference in fluorescence of His-tag fluorescence-labeled RCT before (0 s) and 1.5 s after induction of a laser-induced thermal gradient (normalized fluorescence, *F*_*n*_) increased with titration of ATP and ADP at pH 7.0 and pH 8.0. Shown is the difference in *F*_*n*_ (Δ*F*_*n*_) relative to the signal of ligand-free RCT as determined by the 1:1 model fit using the PALMIST software of representative experiments with n= 3 technical replicates ± SD. No ATP response was observed for RCT_G65A_, while ADP was bound at pH 7.0 but not at pH 8.0. **b**, Addition of ATP or ADP increases the thermal stability of RCT_WT_ at pH 7.0 and even more at pH 8.0. Neither ADP nor ATP changed thermal stability of RCT_G65A_ at pH 8.0. Melting temperatures (T_M_) were determined by differential scanning fluorimetry using protein intrinsic fluorescence as a read out for protein denaturation (Supplemental Fig. 7a). ΔT_M_ was derived from subtracting single measurements from the average T_M_ of ligand-free RCT_WT_. Error bars = SD.; n = 3. Different lowercase letters above bars indicate significant statistical differences between conditions with *P* < 0.05 as calculated by ANOVA and Holm-Sidak pairwise multiple comparison. **c**, Near-UV circular dichroism spectroscopy was used to determine tertiary structure (measured as ellipticity, Θ in mdeg) of RCTWT at pH 7.0 and 8.0. Without the addition of nucleotides, pH had little effect on Θ of RCT_WT_. Addition of ATP and ADP lead to pH-dependent differences in ellipticity as compared to ligand-free RCT_WT_ (ΔΘ).

The function of the RCT may either respond to stromal ATP levels alone or, alternatively, to the phosphorylation potential, which is set by the ratio of ATP and ADP. Thus, we investigated whether the RCT was able to bind ADP. *In silico* docking experiments supported ADP to favor the same RCT binding site as ATP (Supplemental Fig. 6c). MST analysis showed that ADP binding curves for RCTWT resembled at both pH values the ATP binding curve from pH 7.0 (Fig. 6a). For RCT_G65A_ they revealed a low response to ADP at pH 8.0, but a similar response to RCT_WT_ at pH 7.0.

### Binding of ATP induces strong conformational changes

We then investigated the effects of ATP- and ADP-binding on protein conformation. We utilized protein intrinsic fluorescence changes in response to thermal denaturation and determined the change in melting temperature (ΔT_M_). Addition of ADP or ATP to the RCT yielded a positive ΔT_M_ at both pH values, with ATP increasing thermal stability much more than ADP (Fig. 6b, Supplemental Fig. 7a). In line with the absence of binding determined by MST, the T_M_ of RCT_G65A_ at pH 8.0 was not affected by the addition of either of the two nucleotides (Fig. 6b, Supplemental Fig. 7a). To verify that changes in thermostability relate to a conformational rearrangement of the RCT, we measured near-UV circular dichroism (CD) spectra. To resolve differences in protein structure, we used a serial two-cuvette system that allowed the exact subtraction of the ATP contribution to the recorded near-UV CD spectra (see Supplemental Fig 7c). The CD spectrum was very similar for RCTWT at the two different pH values (Fig. 6c). When RCT_WT_ was incubated with ATP and ADP, ellipticity changed at similar wavelengths, but differences were much stronger with ATP than with ADP. When experiments were performed at pH 7.0, differences in ellipticity were particularly evident in the shorter wavelength ranges from 260 to 270 nm, while at pH 8.0, differences were strongest around 280 nm (Fig. 6c). Addition of ATP to the RCT_G65A_ did not have any discernable effects on the near-UV CD spectra, as was expected from MST and thermostability measurements (Supplemental Fig. 7c).

Together, the data show that binding of ADP and ATP changes the conformation of RCT_WT_ in a pH dependent manner. The data are consistent with the scenario that these conformational changes are favored by binding of ATP rather than ADP. The competitive binding of both species and different effects on the stabilization of a certain conformation provides a potential mechanism, by which KEA3 senses the stromal phosphorylation potential and reacts in response.

### NADP^+^ and NADPH induce additional structural changes via a different binding site

Next, we asked whether the RCT also binds NADPH and NADP^+^. *In silico* docking analyses showed NAD species to preferentially bind to a different site than ATP and ADP (Fig. 7a, Supplemental Fig. 8a-b). Similar to the simulations with ATP, NADP^+^ and NADPH led to a decrease in RCT_WT_ radius, albeit with less distinct maxima (Fig. 7b). Interestingly, most secondary structure differences between free RCT and ligand-bound RCT were independent of the added ligand, despite the presence of ATP, ADP and NADPH, NADP+ in two distinctly different binding pockets during simulation (Supplemental Fig. 8d). Both NADP^+^ and NADPH additionally induced specific differences in secondary structure that were distinct from ATP and the other NAD species. While NADPH led to an increase in α-helical structures around the glycine-rich stretch of RF2, NADP^+^ led to the conversion of coils into the extension of a ß-sheet between AA 153-155 (Fig. 7c, Supplemental Fig. 8d, Supplemental Fig. 5a).

**Fig. 7.**
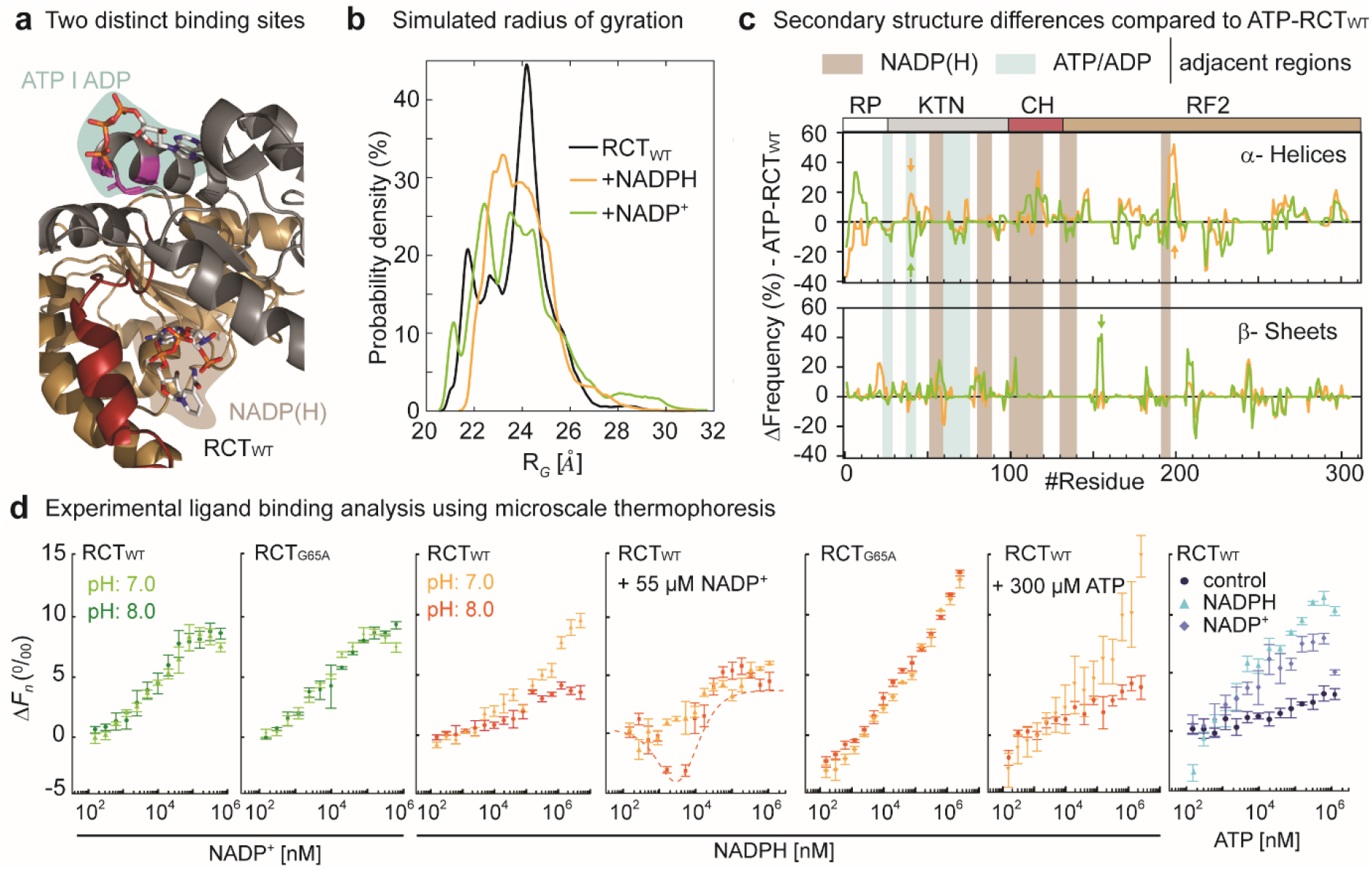
NADP^+^ and NADPH bind at a second nucleotide binding site and induce both ATP-similar and distinct conformational changes during simulation. **a**, Model of the RCT with binding sites for ATP, ADP (blue) and NADP(H) (brown) as identified by *in-silico* docking. **b**, The probability density of radii of gyration was calculated from MD simulations. The analysis shows that both NADPH and NADP^+^ lead to a decrease in RCT radius. **c**, Frequencies of α-helices and ß-sheets along the RCT protein sequence after running MD simulations with NADPH or NADP^+^ subtracted from those with ATP (ΔRCT_WT_+ATP). The ATP, NADP(H)-adjacent regions are shown in light blue or brown respectively. Frequencies are calculated from n = 10 simulations. Arrows indicate simulated secondary structure differences specifically induced by NADPH (orange) or NADP^+^ (green) as compared to ATP. Recombinant part, RP; connecting helix, CH; Rossmann-fold 2, RF2. **d**, Microscale thermophoresis (MST) analysis of RCT_WT_ and RCT_G65A_ reveal very similar patterns of normalized fluorescence changes (Δ*F*_*n*_) for NADP+ titration at pH 7.0 and pH 8.0. NADPH binding in the presence of NADP+ suggests that affinity of RCT_WT_ for NADPH is higher than for NADP+ at pH 8.0. Saturation with both NADPH and NADP+ increases Δ*F*_*n*_ induced by titrating ATP (NADPH more than NADP+), suggesting that binding of either nucleotide changes physical properties of RCT_WT_ synergistically with ATP. Shown is the difference in fluorescence between before and 1.5 s after induction of the laser-induced thermal gradient relative to the signal of unbound RCT (Δ*F*_*n*_) as determined by the PALMIST software of representative experiments with n= 3 technical replicates ± s.d.

By performing MST analysis, we received experimental support for a second nucleotide binding pocket (Fig. 7d). Both the RCTG65A with a mutation in the ATP, ADP binding site and RCTWT showed identical MST traces when titrated with NADP^+^ at both tested pH values, supporting that NADP^+^ binding does not occur at the KTN-nucleotide binding site. An MST competition analyses using NADP^+^ saturated RCTWT and titrating NADPH revealed normalized fluorescence (ΔFn) to first drop and then rise again at pH 8.0. This biphasic behavior can be explained by the following: (i) NADPH displaces NADP^+^ at low concentrations from the competitive binding site, which leads to a change in conformation and thus a decrease in MST fluorescence, and (ii) NADPH at higher concentrations binds the second nucleotide binding site located in the KTN domain, which then increases MST fluorescence. This explanation postulates that at least at pH 8.0, the RCT has a higher affinity for NADPH than for NADP^+^. NADPH titration of ATP saturated RCTWT and vice versa revealed that ATP saturation has no major effect, but that both NADPH and NADP^+^ increase the MST response during ATP titration, with NADPH yielding the highest ΔFn values. Differently to ATP and ADP, binding of NADP^+^ or NADPH did not cause any changes in RCT thermostability (Supplemental Fig. 9), supporting that effects of both types of nucleotides on protein structure are different.

Together, the results are in line with a high affinity binding site for NADP^+^ and NADPH that is distinct from the binding site for ATP and ADP of the KTN domain. Both, MD simulation as well as experimental MST analysis suggest that binding of NADP^+^ and NADPH to this NADP-specific site of RCT has distinct consequences for protein conformation, which are additive to those induced by ATP or ADP at the KTN binding site.

### Activation of KEA3_G531A_ is delayed during high to low light transitions

Finally, we asked how the G>A substitution of the first glycine in the KTN glycine-rich stretch affects KEA3 function *in planta*. We generated plants stably expressing the respective KEA3_G531A_ version as a C-terminal GFP fusion from the native *KEA3* promoter. Visually, these plants were indistinguishable from WT, *kea3-1* and the *KEA3-GFP* in *kea3-1* (*KEA3*) plants (Fig. 8a). Two lines were selected, L5 accumulating *KEA3*_*G531A*_ at WT and L7 at *KEA3*-levels (Fig. 8b). Plants were exposed to changes in light intensity including a transition from high light to low light (Supplemental Fig. 10 a). KEA3 activity is repressed during high light and activated directly after a transition to low light to rapidly downregulate NPQ (Fig. 2c, Fig. 8c). During steady state high light, NPQ was very similar between lines with a significant increase only in *kea3* as compared to the *KEA3* line with twice the amount of KEA3 protein than WT (Fig. 8d). After transition to low light, both lines expressing *KEA3*_*G531A*_ exhibited much higher NPQ than WT and *KEA3* (Fig. 8d). This suggests that KEA3 of this mutated version is not activated to the same degree as in WT, likely because *KEA3G531A* is unable to sense the activating signal and induce structural changes required for full activation. To correlate changes in KEA3 activity with ATP levels *in situ*, we used the stromal targeted MgATP^2-^ sensor ATeam-1.03nD/nA ^19-21^ during light fluctuations (Supplemental Fig. 10b). The FRET response was very minor when compared to the potential response range of the sensor and was within the range by which the background signal responded to the transitions between low and high light. Yet, we observed a steady increase in ATeam FRET intensity during low light phases, when the background signal was stable. The FRET increase was reversed after switch to darkness, consistently with a recent observation^19^.

**Fig. 8.**
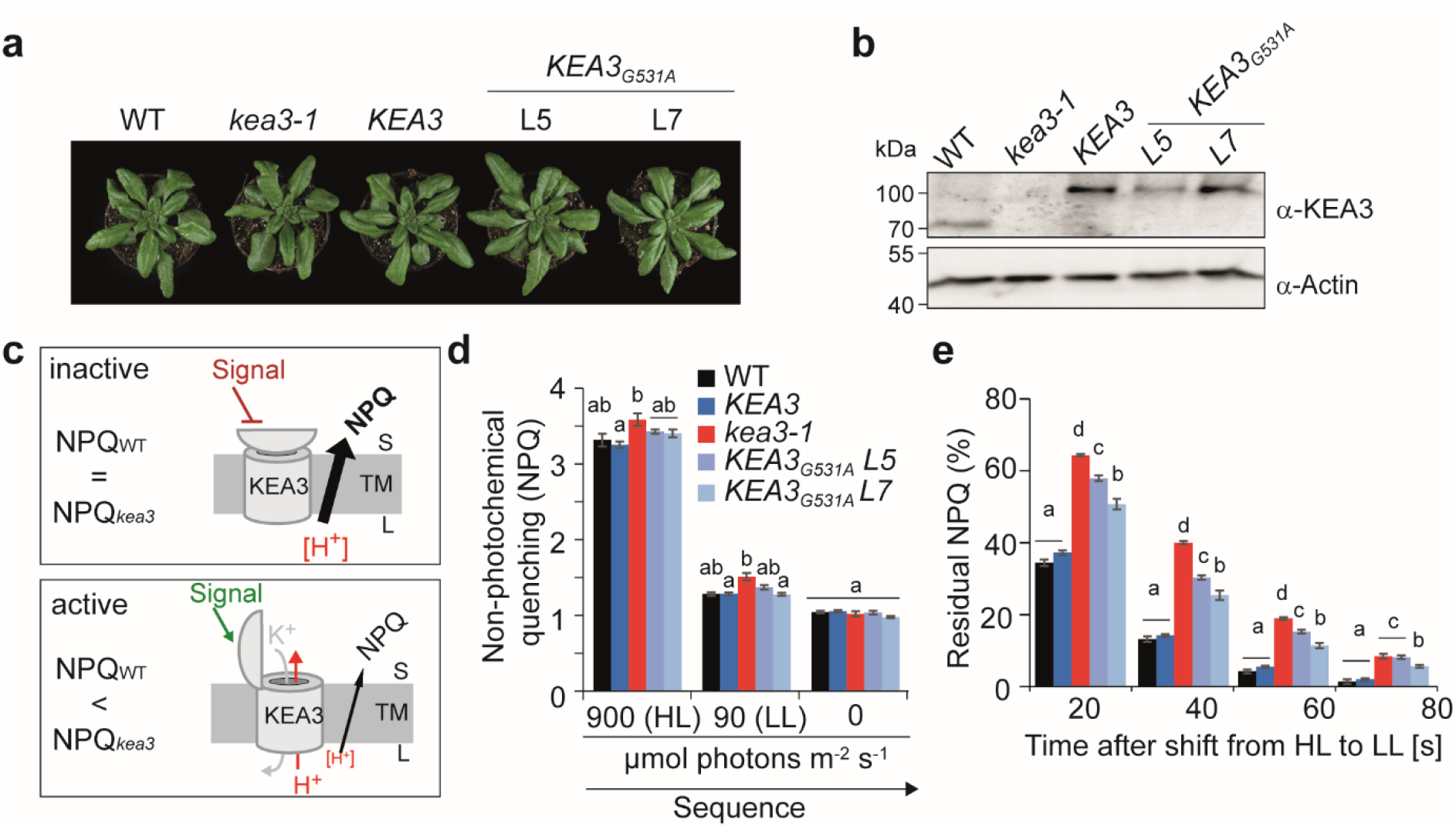
Substitution of KEA3 G531 with A impairs activation after transition from high to low light. **a**, Picture of five-week-old WT, *kea3-1, KEA3* (*KEA3-GFP* in *kea3-1*) and *KEA3*_*G531A*_ (*KEA3*_*G531A*_*-GFP* in *kea3-1*) lines L5 and L7. **b**, KEA3 protein level in the different lines as determined via immuno-blot with the KEA3 antibody and Actin as the loading control. **c**, Model depicting how regulation of KEA3 activity influences non-photochemical quenching (NPQ). **d**, The different lines as described in a and b were exposed to an alternating light regime of 5 min low light (LL, 90 µmol photons m^-2^ s^-1^), 5 min high light (HL, 900 µmol photons m^-2^ s^-1^), 5 min LL and 5 min darkness and non-photochemical quenching (NPQ) was determined from chlorophyll a fluorescence analysis. Values are shown from the end of each different light treatment. **e**, NPQ relaxation was determined after the HL to LL shift and is presented as % of fast relaxing NPQ remaining after the indicated times in LL. **d-e**, Different lowercase letters above bars indicate significant statistical differences between genotypes at the given timepoint with *P* < 0.05 as calculated by ANOVA and Holm-Sidak pairwise multiple comparison. Error bars = s.e.; n = 3 (WT, *kea3-1*), n = 12 (KEA3, *KEA3*_*G531A*_ L5 and L7).

## Discussion

In the current work, we show that the RCT resides in the chloroplast stroma and thus is exposed to changes in this specific molecular environment. We demonstrate that stromal pH undergoes light intensity dependent changes and that KEA3 regulation coincides with large alterations in stromal pH that occur upon transitions in light intensity. Our results show that a rapid increase in light intensity induces simultaneous opposite changes in the pH of lumen and stroma, generating a jump in ΔpH across the thylakoid membrane. Here, RCT-mediated inhibition is needed to counteract the dissipation of ΔpH by KEA3.

Because the RCT contains a KTN domain with conserved nucleotide binding function, regulation of KEA3 activity was proposed to be triggered by nucleotides^6,7^. Previous reports pointed to the phosphorylation potential or single adenylate species as potential regulators^10,22^, while a computational study correlated KEA3 regulation with the redox potential of the stroma^23^. We show that the KTN nucleotide binding site of the RCT binds ATP and ADP. While pH alone has little effect on protein conformation, structural changes in response to ATP and ADP binding are pH dependent. These structural changes are more strongly induced by ATP than by ADP, suggesting that binding of ATP more effectively stabilizes the change in conformation. MD simulations suggest ATP-dependent differences in conformation to involve the glycine-rich stretch and adjacent regions as well as CH and RF2. Both, computational and experimental approaches, point to the RCT decreasing its radius in response to ATP binding.

Our study reveals that the RCT binds NADPH and NADP^+^ at a second binding site. MD simulations and MST results suggest that binding of NADPH has different consequences for RCT conformation than NADP^+^. While NADPH may also bind to the KTN nucleotide binding site at higher concentrations, it does not affect the thermostability of RCT, which is in contrast to ATP and ADP. This suggests that if bound to the KTN binding site, NADPH does not induce the same structural changes as ATP or ADP. Saturation of the RCT with NADPH and MST analysis with ATP suggests an additive effect of both nucleotides on protein conformation.

By exchanging the first glycine of the glycine-rich stretch with alanine, we generated an RCT version that is unable to bind ATP at either of the two tested physiological pH values and ADP at pH 8.0. During MD simulations this mutated version adopted a similar conformation as the RCT in complex with ATP. Introducing this mutation into full-length KEA3 *in planta*, revealed that the first glycine of the glycine-rich stretch is crucial for activation of KEA3 in response to a shift from high to low light.

Together, our data support a model in which the energy status of the chloroplast is measured by the KEA3 RCT via three different signals, the phosphorylation potential (ATP/ADP), the redox potential (NADPH/NADP^+^) and stromal pH, which act synergistically on protein conformation. By careful measurement of stromal pH, estimation of stromal MgATP^2-^ and literature-derived information on stromal redox potential of the NADP pool^24-26^, we propose that during high light phases, a combination of high pH and redox potential, together with elevated phosphorylation potential results in deactivation of KEA3 (Fig. 9). This deactivation involves conformational changes of the RCT, which may be transduced to additional changes in the transport domain.

**Fig. 9.**
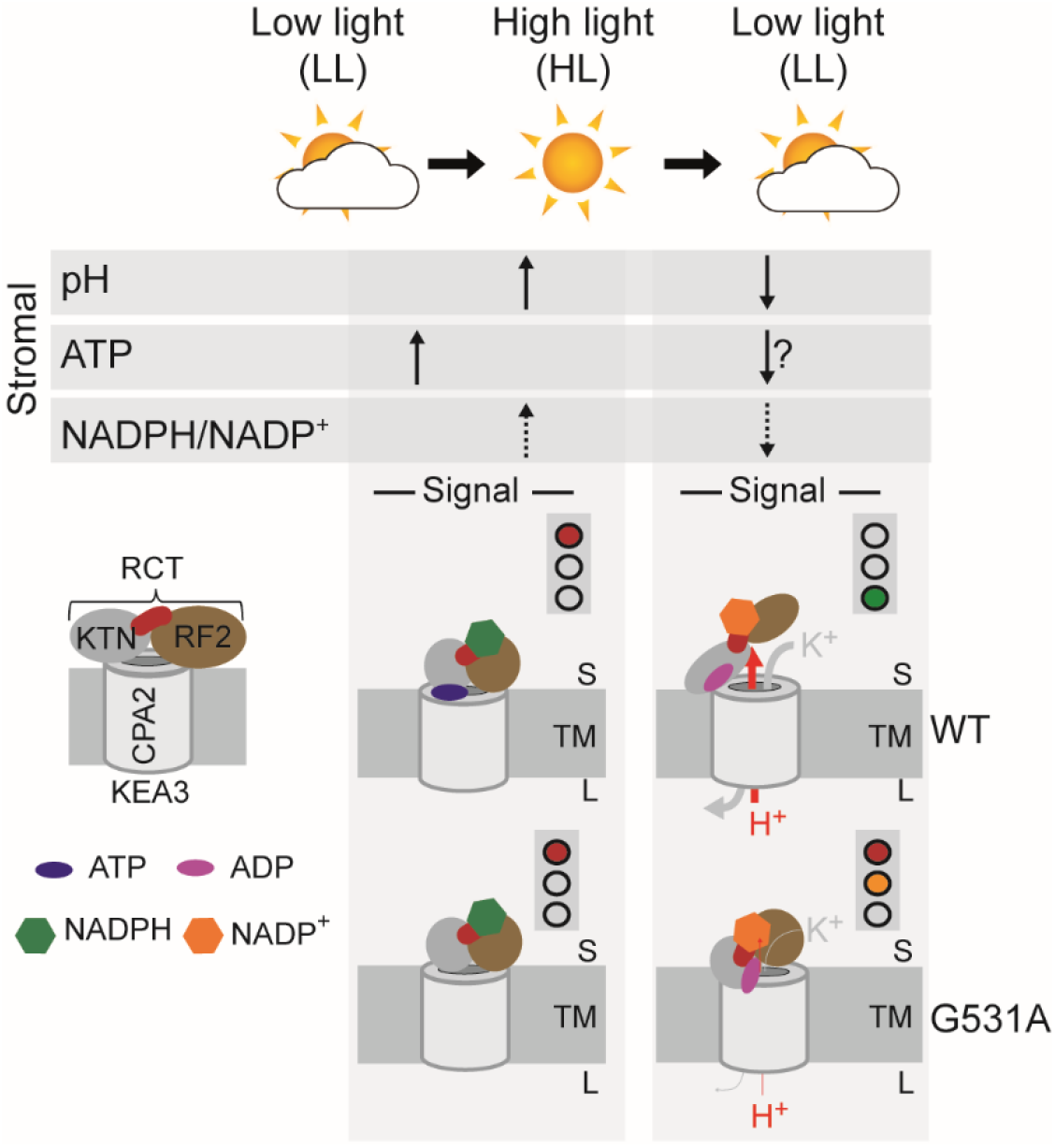
Model of KEA3 regulation in response to stromal signals. Model summarizing changes in stromal factors during changes in light intensity and their signaling effect on KEA3 activity. Stromal pH and ATP were determined in this study with leaving a question mark for ATP after the high to low light transition, because of low signal to noise ratio. NADPH/NADP+ is inferred from published literature. KEA3 is inactivated after transition from low to high light and this coincides with an increase in stromal pH, an already increased level of ATP, which slowly increases during low light illumination and an increase in the NADPH/NADP^+^ ratio. KEA3 is activated during a high to low light transition, which coincides with a decrease in stromal pH, possibly a decrease in ATP and likely a decrease in NADPH/NADP^+^. All three stromal factors act synergistically on the conformation of the KEA3 RCT, which we hypothesize results in the change of activity.

Taken together, the regulation of KEA3 activity appears to be highly complex provoking the question as to why such complexity is needed. We hypothesize that the synergistic action of the different signals allows a for plastic response of KEA3 and thus qE to metabolic requirements in a highly flexible manner. Whether changes in RCT signal responses may be exploited to enhance qE dynamics, remains to be determined.

In summary, we present a regulatory feedback loop by which the energy state of the chloroplast, in form of the phosphorylation and redox potential in combination with stromal pH, adjusts qE dynamics via KEA3 mediated proton antiport. These findings have broad implications for our basic understanding of plant energy metabolism, metabolic engineering of plants, and enhancing photosynthesis in changing environments.

## Material and Methods

### Self-assembly GFP analysis

For cloning of the different constructs see Supplemental Table 1. The plasmids encoding saGFP1 and saGFP2 variants were co-transformed into Arabidopsis protoplasts according to Gerdes et al, 2009^27^. Fluorescence of protoplasts was captured using a Leica TCS SP5II inverted confocal laser scanning microscope (Leica Wetzlar, Germany) with a 40x water immersion objective. For excitation a wavelength of 488 nm was used and eGFP and chlorophyll fluorescence were detected between 495 and 565 nm, and 580 and 670 nm, respectively.

### Plant material and growth conditions

Information on plant material can be found in Supplemental Information. Wild type (Col-0) and transgenic Arabidopsis lines were sown on soil, under long-day controlled conditions (LD, 16h day at 20 ^°^C/8h night at 6 ^°^C with relative humidity of 60% and 75% at day and at night respectively), at 250 µmol photons m^-2^ s^-1^ for seven days. Afterwards, plants were moved to a long day condition chamber with 120 – 150 µmol photons m^-2^ s^-1^ and 16 ^°^C temperature at night. Plants were pricked into the individual pots after 2 weeks. All experiments were performed with 4 – 5 weeks old plants.

### Nucleotide affinity purification from isolated thylakoid membranes

Thylakoid membranes according to 400 µg chlorophyll were solubilized with 1% ß-DM (n-dodecyl-ß-D-maltoside) in IP buffer at pH 8.0 (50 mM HEPES/KOH pH 8.0.0, 330 mM sorbitol, 150 mM NaCl, 1 mM PMSF, and protease inhibitor cocktail) or pH 6.8 (50 mM MES/KOH pH 6.8, 330 mM sorbitol, 150 mM NaCl, 1 mM PMSF, and protease inhibitor cocktail) and incubated with nucleotide-linked agarose beads (AMP and ATP affinity test kits, Jena Biosciences, Jena, Germany) overnight on the wheel at 4^°^C at a final concentration of 0.2% ß-DM. Beads were washed 5x with the same buffer (3x + 0.2% ß-DM, then 2x without) before proteins were eluted with SDS.

### Protein analysis

Thylakoids and total leaf proteins were isolated as previously described^6,28^. SDS-PAGE and immunoblot analyses were performed according to Uflewski et al., 2021^9^. Freshly isolated thylakoid membranes were treated with chaotropic salts according to Karnauchov et al., 1997^29^. Antibodies specific for the C-terminus of KEA3^6^, GFP (Chromotek), PetA and Actin (Agrisera) were used for protein detection.

### Sensing of stromal pH and MgATP2-by chloroplast targeted cpYFP and ATeam

Stromal dynamics during light transition were monitored using the TCS SP8 microscope (Leica, Wetzlar, Germany) equipped with a x20 lens (HC PL APO 20x/0.75 CS2, dry immersion). A customized lightbox was manually switched between two light intensities (high light: 900 and low light: 90 µmol photons m^-2^ s^-1^). Light was provided by a 620-645 nm LED (LXM2-PD01-0060, Lumileds, Amsterdam, NL) flexibly mounted to the dish holder of the motorized stage. Confocal imaging was performed using discs of true leaves of 4-to 5-week-old *Arabidopsis thaliana* plants. The leaf discs were mounted between two cover slips and the LED was directly attached from the top side. For pH measurements GFP or YFP fluorescence was excited at 488 nm, while emission was collected at 498 – 548 nm. The ATeam sensor for MgATP^2-^ was excited at 458 nm and emission was recorded at 470 – 507 nm for CFP and 521 – 531 nm for YFP. Chlorophyll fluorescence was collected simultaneously at 650 – 700 nm. Resolution was set to 512×512 pixel and 0.6475 s frame time, and pinhole to 142 µm. Plants were dark-acclimated for at least 30 min before starting the measurements. Fluorescence levels were collected from the entire microscopy frame and baseline corrected using the last minute of each dark-phase (before and after the light treatment). To ensure that pH changes were measured in photosynthetic active plastids, the focal plane was placed in the mesophyll chloroplasts of the subepidermal adaxial leaf side, the palisade parenchyma.

### Molecular modelling

For molecular modelling of the experimentally examined KEA3 RCT, the software MODELLER version 9.25^30^ was utilized with the crystal structure of the C-terminal domain of the KefC-KefF complex from *Escherichia coli* (PDB ID 3eyw, https://www.rcsb.org) serving as the template. The accuracy of the predicted models was evaluated by the software’s molpdf, DOPE, and GA341 scores and the stereo-chemical properties were validated by the online tool MolProbity^31^. The investigated RCT_G65A_ mutant was built based on the predicted WT structure. The pH depended protonation states of amino acid residues were computed by the H++ online tool^32^.

### Ligand docking

Protein-ligand docking studies were performed by SwissDock^34^ with ligand structures from the ZINC database^35^. The resulting protein-ligand complexes were evaluated by the SwissDock fitness value^36^. Docked models with the best (lowest) scores were selected for MD studies.

### Molecular dynamics simulation

All MD simulations were performed using the Gromacs molecular dynamics simulation engine^37^ version 2020.5/2020.6 and CHARMM36 (C36) force field^38^. Each protein and protein-ligand model was centered in a dodecahedral simulation box with a minimum edge length of 20 Å. Further information can be found in the supplemental material.

The progress of each resulting MD trajectory was evaluated by the root mean square deviation (RMSD) of protein backbone atoms and the radius of gyration (R_G_) using the Gromacs rms or gyrate tools, respectively. The secondary structure was assigned using the do_dssp tool based on the DSSP algorithm^39,40^. The distance between each RCT isoform residue and the nucleotide was determined using Gromacs mindist algorithm and a contact was defined if any two atoms of each interaction groups were within a cutoff of 7 Å.

### Heterologous expression and protein purification

The RCTG65A point mutation was introduced in the pET28a (+) vector (Novagen) containing the regulatory C-terminus of KEA3 (RCT: KEA3_AA495-776_^41^) by using specific primers as detailed in Supplemental table 2. Protein expression of RCT and RCT_G65A_ and purification were performed as described previously for RCT^42^ with the following modifications: protein containing bacterial pellets were disrupted using the EmlusiFlex C-3 (AVESTIN, Inc.) and the soluble fraction was obtained by centrifugation at 18,000 *g* and subsequent 0.2 µM pore size filtration. Recombinant proteins were purified by using Ni^2+^ NTA-agarose and elution with 300 mM imidazole, treated with 2 mM EDTA and separated by size exclusion chromatography to remove residual Ni^2+^ ions and protein aggregates. Subsequently, the RCT fraction was dialyzed against phosphate-buffered saline (PBS) at pH 8.0 to remove the imidazole and concentrated by Amicon centrifugation, both using membranes with 10 kDa cutoff.

### Microscale thermophoresis

Protein sample preparation and measurements were performed according to the protocol from Monolith NT (NanoTemper Technologies GmbH, Munich, Germany), employing quality controls as outlined by Sedivy 2021^43^. The N-terminal His-tag was labelled with the RED-tris-NTA kit (NanoTemper Technologies GmbH, Munich, Germany). After 10 minutes incubation on ice, 50 nM of protein sample in PBS together with indicated concentrations of nucleotides were loaded into standard Monolith NT.115 capillaries (NanoTemper Technologies GmbH, Munich, Germany). MST measurement was performed using the Monolith NT.115 instrument (NanoTemper Technologies GmbH, Munich, Germany) at ambient temperature of 23 ^°^C. Instrument parameters were adjusted to 80 % LED and 60 % MST power. The difference in normalized fluorescence was calculated between the initial fluorescence and 1.5 s after the onset of the laser. 10 mM nucleotide stocks were prepared freshly in PBS pH 8.0 directly before the experiment and the pH was checked. Data of three replicates were analyzed and fitted using the PALMIST software^44^.

### Protein structure and stability analyses by differential scanning fluorimetry and circular dichroism spectroscopy

The purified protein in PBS at a concentration of ∼16 µM was loaded into capillaries (nanoDSF grade standard, NanoTemper GmbH, Munich, Germany) and a melting curve was determined by using the nanoDSF Prometheus NT.48 instrument (NanoTemper GmbH, Munich, Germany) set to an LED power of 60% to 80%, a temperature ramp ranging from 20-75 ^°^C and data collection at thermal intervals of 1.5 ^°^C. Calculation of melting temperature inflection points was performed using PR.Therm Control software (NanoTemper GmbH, Munich, Germany).

For near-UV circular dichroism (CD) spectroscopy, 100 µM protein in PBS at pH 8.0 with or without ATP was loaded into a quartz cuvette with 2 mm path length (Hellma GmbH, Germany). Measurements were performed at 16 ^°^C using a Jasco J-715 spectropolarimeter (Jasco Deutschland GmbH, Pfungstadt, Germany) equipped with a Peltier thermostat-controlled cell holder, simultaneously with an in-line localized 2 mm path length reference cuvette containing PBS buffer or ATP dissolved in PBS buffer at concentrations corresponding to those in the protein samples. Further experimental information can be found in the supplemental material.

### Determination of apparent masses and hydrodynamic radii by SEC and light scattering

Purified RCT was dissolved in PBS pH 8.0 and chromatographed on a Superdex 75 Increase 3.2/300 column (Cytiva, Marlborough, MA, USA) using an NGC chromatography system (BioRad Laboratories GmbH, Feldkirchen, Germany). SEC was performed in PBS pH 8.0 at 8 °C with a flow rate of 0.04 ml/min. The retention volume of the main protein peak was determined and protein mass and hydrodynamic radius were calculated based on the retention volumes of a set of reference proteins from the Cytiva Gel Filtration LMW Calibration Kit (Cytiva, Marlborough, MA, USA).

Simultaneous static (SLS) and dynamic light scattering (DLS) experiments were performed as previously described in Gats et al., 2017^45^. For further experimental details, see the supplemental material. The protein concentration was measured photometrically using a specific absorbance A (1 cm path length, 1 mg/ml) of 0.387 at 280 nm for samples without ATP and 0.128 at 293 nm in presence of ATP. To correct for solvent effects concentration dependent data of apparent masses and hydrodynamic radii of RCT in presence and absence of ATP were extrapolated to zero protein concentration yielding the mass and hydrodynamic radius of the protein.

### Chlorophyll *a* fluorescence measurement

Chlorophyll *a* fluorescence was recorded of dark-acclimated plants using the Imaging PAM (Waltz GmbH, Effeltrich, Germany). Saturation light pulses (setting: 10) were applied after dark acclimation (for Fm) and illumination (for Fm’). The non-photochemical quenching (NPQ) at a given time point during the light treatments was calculated as (Fm − Fm′)/Fm′^46^.

## Acknowledgement

U.A. received funding from the DFG (AR 808/5-1) and the Max Planck Society. We thank Markus Miettinen for help with the MD simulations.

## References

1 Kaiser, E., Morales, A. & Harbinson, J. Fluctuating Light Takes Crop Photosynthesis on a Rollercoaster Ride Plant Physiology 176, 977–989, doi:10.1104/pp.17.01250 %J Plant Physiology (2017).

2 Roden, J. S. & Pearcy, R. W. Effect of leaf flutter on the light environment of poplars. Oecologia 93, 201–207, doi:10.1007/BF00317672 (1993).

3 Külheim, C., Agren, J. & Jansson, S. Rapid regulation of light harvesting and plant fitness in the field. Science (New York, N.Y.) 297, 91–93, doi:10.1126/science.1072359 (2002).

4 De Souza, A. P. et al. Soybean photosynthesis and crop yield are improved by accelerating recovery from photoprotection. 377, 851–854, doi:doi:10.1126/science.adc9831 (2022).

5 Kromdijk, J. et al. Improving photosynthesis and crop productivity by accelerating recovery from photoprotection. 354, 857–861, doi:doi:10.1126/science.aai8878 (2016).

6 Armbruster, U. et al. Ion antiport accelerates photosynthetic acclimation in fluctuating light environments. Nat Commun 5, 5439, doi:10.1038/ncomms6439 (2014).

7 Armbruster, U. et al. Regulation and Levels of the Thylakoid K+/H+ Antiporter KEA3 Shape the Dynamic Response of Photosynthesis in Fluctuating Light. Plant Cell Physiol 57, 1557–1567, doi:10.1093/pcp/pcw085 (2016).

8 Kunz, H. H. et al. Plastidial transporters KEA1, -2, and -3 are essential for chloroplast osmoregulation, integrity, and pH regulation in Arabidopsis. Proc Natl Acad Sci U S A 111, 7480–7485, doi:10.1073/pnas.1323899111 (2014).

9 Uflewski, M. et al. Functional characterization of proton antiport regulation in the thylakoid membrane. Plant Physiol, doi:10.1093/plphys/kiab135 (2021).

10 von Bismarck, T. et al. Light acclimation interacts with thylakoid ion transport to govern the dynamics of photosynthesis. Research Square, doi:10.21203/rs.3.rs-948381/v1 (2021).

11 Wang, C. et al. Fine-tuned regulation of the K(+) /H(+) antiporter KEA3 is required to optimize photosynthesis during induction. Plant J 89, 540–553, doi:10.1111/tpj.13405 (2017).

12 Cao, Y. et al. Gating of the TrkH ion channel by its associated RCK protein TrkA. Nature 496, 317–322, doi:10.1038/nature12056 (2013).

13 Kröning, N. et al. ATP binding to the KTN/RCK subunit KtrA from the K+ -uptake system KtrAB of Vibrio alginolyticus: its role in the formation of the KtrAB complex and its requirement in vivo. J Biol Chem 282, 14018–14027, doi:10.1074/jbc.M609084200 (2007).

14 Roosild, T. P., Miller, S., Booth, I. R. & Choe, S. A mechanism of regulating transmembrane potassium flux through a ligand-mediated conformational switch. Cell 109, 781–791, doi:10.1016/s0092-8674(02)00768-7 (2002).

15 Schrecker, M., Wunnicke, D. & Hänelt, I. How RCK domains regulate gating of K+ channels %J Biological Chemistry. 400, 1303–1322, doi:doi:10.1515/hsz-2019-0153 (2019).

16 Pliotas, C. et al. Adenosine Monophosphate Binding Stabilizes the KTN Domain of the Shewanella denitrificans Kef Potassium Efflux System. Biochemistry 56, 4219–4234, doi:10.1021/acs.biochem.7b00300 (2017).

17 Armbruster, U., Correa Galvis, V., Kunz, H. H. & Strand, D. D. The regulation of the chloroplast proton motive force plays a key role for photosynthesis in fluctuating light. Curr Opin Plant Biol 37, 56–62, doi:10.1016/j.pbi.2017.03.012 (2017).

18 Wiesemann, K., Groß, L. E., Sommer, M., Schleiff, E. & Sommer, M. S. in Membrane Biogenesis: Methods and Protocols (eds Doron Rapaport & Johannes M. Herrmann) 131–144 (Humana Press, 2013).

19 Elsässer, M. et al. Photosynthetic activity triggers pH and NAD redox signatures across different plant cell compartments. 2020.2010.2031.363051, doi:10.1101/2020.10.31.363051 %J bioRxiv (2020).

20 De Col, V. et al. ATP sensing in living plant cells reveals tissue gradients and stress dynamics of energy physiology. eLife 6, doi:10.7554/eLife.26770 (2017).

21 Voon, C. P. et al. ATP compartmentation in plastids and cytosol of <em>Arabidopsis thaliana</em> revealed by fluorescent protein sensing. 115, E10778–E10787, doi:10.1073/pnas.1711497115 %J Proceedings of the National Academy of Sciences (2018).

22 Correa Galvis, V. et al. H(+) Transport by K(+) EXCHANGE ANTIPORTER3 Promotes Photosynthesis and Growth in Chloroplast ATP Synthase Mutants. Plant Physiol 182, 2126–2142, doi:10.1104/pp.19.01561 (2020).

23 Li, M. et al. Impact of ion fluxes across thylakoid membranes on photosynthetic electron transport and photoprotection. Nature plants 7, 979–988, doi:10.1038/s41477-021-00947-5 (2021).

24 Heber, U. W. & Santarius, K. A. Compartmentation and reduction of pyridine nucleotides in relation to photosynthesis. Biochimica et Biophysica Acta (BBA) - Biophysics including Photosynthesis 109, 390–408, doi:https://doi.org/10.1016/0926-6585(65)90166-4 (1965).

25 Lim, S.-L. et al. In planta study of photosynthesis and photorespiration using NADPH and NADH/NAD+ fluorescent protein sensors. Nature Communications 11, 3238, doi:10.1038/s41467-020-17056-0 (2020).

26 Smith, E. N., Schwarzländer, M., Ratcliffe, R. G. & Kruger, N. J. Shining a light on NAD- and NADP-based metabolism in plants. Trends Plant Sci 26, 1072–1086, doi:10.1016/j.tplants.2021.06.010 (2021).

27 Gerdes, L. et al. A Second Thylakoid Membrane-localized Alb3/OxaI/YidC Homologue Is Involved in Proper Chloroplast Biogenesis in Arabidopsis thaliana*. Journal of Biological Chemistry 281, 16632–16642, doi:https://doi.org/10.1074/jbc.M513623200 (2006).

28 Armbruster, U. et al. Regulation and Levels of the Thylakoid K+/H+Antiporter KEA3 Shape the Dynamic Response of Photosynthesis in Fluctuating Light. Plant and Cell Physiology, pcw085, doi:10.1093/pcp/pcw085 (2016).

29 Karnauchov, I., Herrmann, R. G. & Klösgen, R. B. Transmembrane topology of the Rieske Fe/S protein of the cytochrome b6/f complex from spinach chloroplasts. 408, 206–210, doi:https://doi.org/10.1016/S0014-5793(97)00427-4 (1997).

30 Sali, A. & Blundell, T. L. Comparative protein modelling by satisfaction of spatial restraints. Journal of molecular biology 234, 779–815, doi:10.1006/jmbi.1993.1626 (1993).

31 Davis, I. W. et al. MolProbity: all-atom contacts and structure validation for proteins and nucleic acids. Nucleic acids research 35, W375–383, doi:10.1093/nar/gkm216 (2007).

32 Anandakrishnan, R., Aguilar, B. & Onufriev, A. V. H++ 3.0: automating pK prediction and the preparation of biomolecular structures for atomistic molecular modeling and simulations. Nucleic acids research 40, W537–541, doi:10.1093/nar/gks375 (2012).

33 van Zundert, G. C. P. et al. The HADDOCK2.2 Web Server: User-Friendly Integrative Modeling of Biomolecular Complexes. Journal of molecular biology 428, 720–725, doi:https://doi.org/10.1016/j.jmb.2015.09.014 (2016).

34 Grosdidier, A., Zoete, V. & Michielin, O. SwissDock, a protein-small molecule docking web service based on EADock DSS. Nucleic acids research 39, W270–277, doi:10.1093/nar/gkr366 (2011).

35 Sterling, T. & Irwin, J. J. ZINC 15--Ligand Discovery for Everyone. J Chem Inf Model 55, 2324–2337, doi:10.1021/acs.jcim.5b00559 (2015).

36 Grosdidier, A., Zoete, V. & Michielin, O. Fast docking using the CHARMM force field with EADock DSS. Journal of computational chemistry 32, 2149–2159, doi:10.1002/jcc.21797 (2011).

37 Abraham, M. J. et al. GROMACS: High performance molecular simulations through multi-level parallelism from laptops to supercomputers. SoftwareX 1-2, 19–25, doi:https://doi.org/10.1016/j.softx.2015.06.001 (2015).

38 Best, R. B. et al. Optimization of the additive CHARMM all-atom protein force field targeting improved sampling of the backbone φ, ψ and side-chain χ(1) and χ(2) dihedral angles. Journal of chemical theory and computation 8, 3257–3273, doi:10.1021/ct300400x (2012).

39 Kabsch, W. & Sander, C. Dictionary of protein secondary structure: pattern recognition of hydrogen-bonded and geometrical features. Biopolymers 22, 2577–2637, doi:10.1002/bip.360221211 (1983).

40 Touw, W. G. et al. A series of PDB-related databanks for everyday needs. Nucleic acids research 43, D364–368, doi:10.1093/nar/gku1028 (2015).

41 Galvis, V. C. et al. H+ transport by K+ exchange antiporter3 promotes photosynthesis and growth in chloroplast ATP synthase mutants. Plant physiology 182, 2126–2143, doi:10.1104/PP.19.01561 (2020).

42 Uflewski, M. et al. Functional characterization of protonantiport regulation in the thylakoid membrane. Plant physiology, 1–21, doi:10.1093/plphys/kiab135 (2021).

43 Sedivy, A. Standard operating procedure for NanoTemper Monolith measurements. European Biophysics Journal 50, 381–387, doi:10.1007/s00249-021-01534-4 (2021).

44 Tso, S. C. et al. Using two-site binding models to analyze microscale thermophoresis data. Analytical Biochemistry, doi:10.1016/j.ab.2017.10.013 (2018).

45 Gast, K. et al. Rapid-Acting and Human Insulins: Hexamer Dissociation Kinetics upon Dilution of the Pharmaceutical Formulation. Pharmaceutical research 34, 2270–2286, doi:10.1007/s11095-017-2233-0 (2017).

46 Baker, N. R. Chlorophyll fluorescence: a probe of photosynthesis in vivo. Annual review of plant biology 59, 89–113, doi:10.1146/annurev.arplant.59.032607.092759 (2008).

