## Supplemental Fig. 1 for "The K^+^ exchange antiporter 3 senses the chloroplast energy status to synchronize photosynthesis"

Supplemental Material & Methods, Supplemental Figures 1-11, Supplemental Tables 1-3, Supplemental References

### Materials and Methods

### Generation of plant material

*KEA3-eGFP* and *KEA3$_{ΔRCT}$-eGFP* were described previously as 2x*KEA3.2-GFP* and 2x*KEA3.3-GFP*[1] and renamed here for simplification. *oeKEA3-eGFP*, *cTP-cpYFP* and *cTP-ATEAM* plants used for confocal microscopy analysis during light fluctuations were published previously in Armbruster et al., 2016[2], Behera et al., 2018[3] and de Col et al., 2017[4], respectively. Plants expressing *KEA3$_{G531A}$*-GFP under the native *KEA3* promoter were generated by site-directed mutagenesis with primers listed in Supplementary Table S1. WT and *kea3-1*[5] over-expressing a chloroplast targeted *RCT* were generated by inserting coding sequences, amplified using primers as specified in Supplementary Table 2, of TSrbcs (from pAVA-TSrbcs-saGFP11N), the RCT (for KEA3 AAs 495-776) and a Myc tag into the *Xba*I and *Sac*I sites of pGWB2[6] downstream of the *35S* promoter. The resulting constructs were transformed into WT and *kea3-1* plants by *Agrobacterium tumefaciens* mediated floral dip[7]. Individual T1 transgenic plants were selected based on their resistance to Basta on soil or to hygromycin on plates and expression of KEA3 or RCT as evaluated by SDS-PAGE analysis and immunodetection using the specific KEA3 antibody.

**Molecular dynamics simulation**

Explicit TIP4P-water[8] solvent box replicates were placed into the simulation system by the tool solvate. Gromacs genion algorithm was used to add sodium ions for the electro-static neutralization of the simulation systems and 0.16 M NaCl mimicking the experimental ionic strength of PBS. Prior to the production runs, the steepest descent algorithm was executed for an energy minimization of the system. This procedure was followed by a solvent equilibration step (NVT) using a Bussi–Donadio–Parrinello thermostat[9] and then an NPT-equilibration performed by the Parrinello-Rahmam algorithm[10]. To avoid premature structural changes, the protein models were kept constrained by the LINCS algorithm[11] during two equilibration processes with simulation times of 1 ns each. Energy minimization and system equilibration steps were considered successful, when they converged to a minimal energy, temperature, pressure and density equilibria over time, respectively. The such equilibrated MD systems were used for further production runs of 1 µs each, which were conducted at a temperature of 300 K with time steps of 2 fs, periodic boundary conditions, a 10 Å spherical cut-off for non-bonded interactions, a force-switch function of 10 Å for van der Waals terms and the Coulomb-type PME (particle-mesh Ewald)[12] was chosen with a 10 Å radius. For all approaches, ten independent repeats were performed.

**Circular dichroism spectroscopy**

The RCT spectra were recorded from 330 nm to 250 nm and the corresponding solvent spectra were subtracted in order to exclude spectral contributions of buffer and ATP. Far-UV spectra were measured in 50 mM phosphate buffer pH 8.0 with 137 mM NaF due to the strong absorption of NaCl contained in PBS and the protein concentration was adjusted with the same buffer to 19.75 µM, 6.47 µM and 3.23 µM and controlled by absorbance measurements at 280 nm using the molar extinction coefficient 12950 $M^{-1} cm^{-1}$. Spectra were recorded from 260 nm to 205 nm for 19.75 µM, from 260 nm to 191 nm for 6.47 µM and from 260 nm to 181 nm for 3.23 µM. Measured far-UV CD spectra were transformed from millidegrees (mdeg) to molar mean residue weight ellipticity ($\Theta_{MRW}$). Spectra from all protein concentration were averaged, asthere were only minor concentration dependent differences. From the far-UV CD spectra the ratios of the secondary structure elements α-helix, β-sheet, turn and coil were estimated by the software CDPro[13] utilizing the three algorithms CONTINLL, CDSSTR and SELCON3 and the reference spectra sets of native globular proteins and denatured proteins.

**Dynamic and static light scattering**

Experiments were performed at a scattering angle of 90° with a custom-built apparatus equipped with a 0.5 W diode pumped continuous-wave laser (Cobolt Samba 532 nm, Cobolt AB, Solna, Sweden), a high quantum yield avalanche photodiode, and an ALV 7002/USB 25 correlator (ALVGmbH, Langen, Germany). Briefly, all measurements were carried out in 3 mm path length microfluorescence cells (105.251-QS, Hellma, Germany) in a Peltier thermostat-controlled cell holder at 15 °C. Prior to the measurements all samples were ultracentrifuged for 30 min at 60,000 g. Apparent hydrodynamic radii were calculated using the Stokes−Einstein equation, $R_S = k_B T/(6\pi\eta D)$, where $k_B$ is the Boltzmann constant, T is the absolute temperature, and η is the solvent viscosity. Apparent molecular masses were calculated by the equation $Mapp = k_{opt} * I_{ex}/c$, where c is the protein concentration, $I_{ex}$ the excess scattering of the protein and $k_{opt}$ is an optical constant depending on physical quantities of the scattering experiment as the scattering angle, wavelength, reference sample, refractive index n of the solution, and refractive index increment (dn/dc) of the protein. We used a refractive index increment (dn/dc) of 0.188 mL/g. The refractive indices of solvents and solutions were determined at 23 °C using an Abbe refractometer, and solvent viscosities were measured using an Ubbelohde-type viscometer (Viscoboy-2, Lauda, Germany).

Supplemental Figures

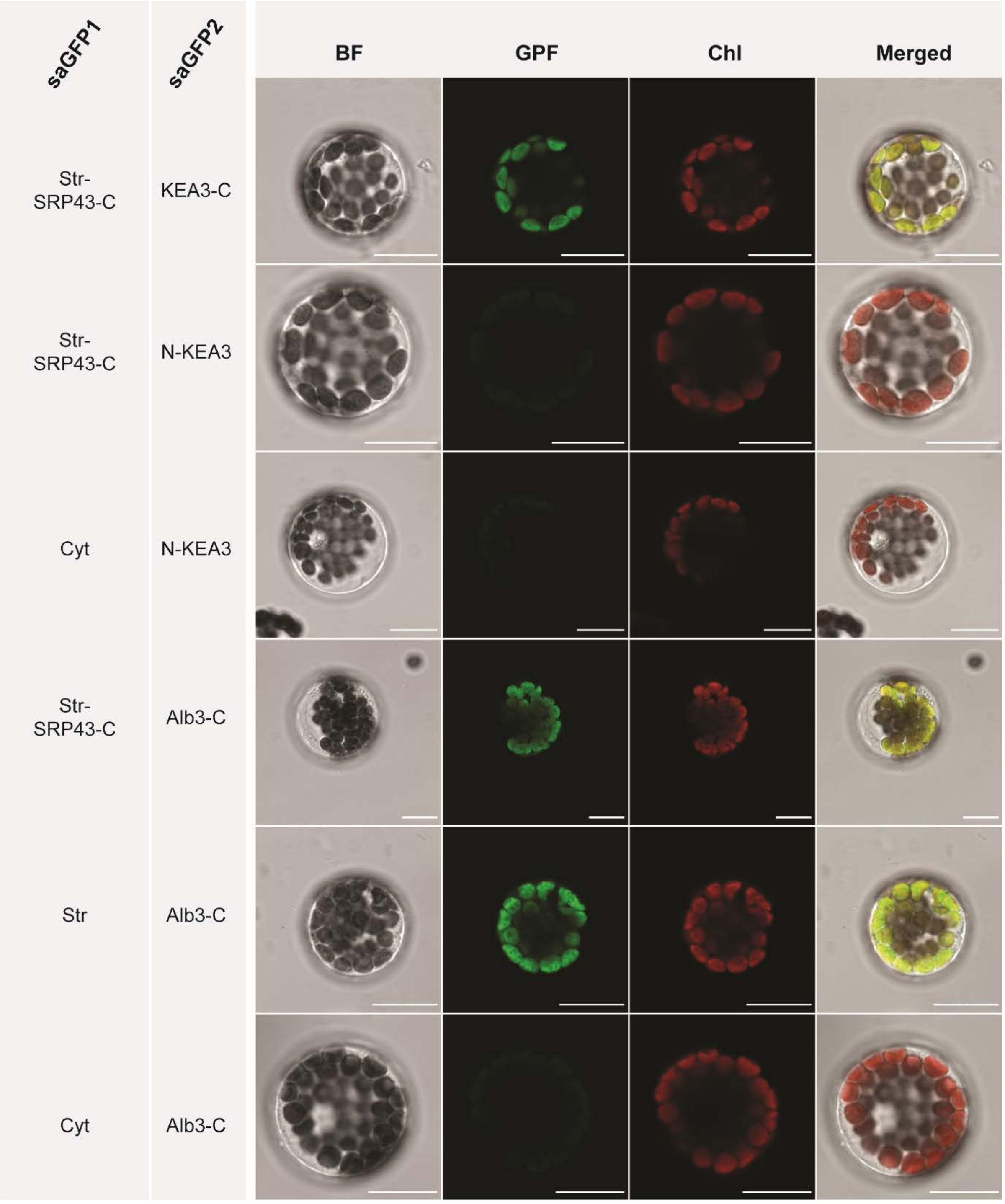


**Supplemental Fig. 1 Controls for the self-assembly GFP analysis**

The larger self-assembly GFP fragment 1 (saGFP1) was either targeted to the stroma by a chloroplast transit peptide (TSrbcs, sequence for chloroplast targeting of Arabidopsis RbcS) or as a fusion with precursor cpSRP43. The smaller saGFP fragment 2 (saGFP2) was fused to the C-terminus of KEA3 (KEA3-C) or Alb3 (Alb3-C). Additionally, it was inserted between the TSrbcs and the mature KEA3 yielding a chloroplast targeted saGFP2-N-KEA3 fusion. Arabidopsis protoplasts were co-transformed with saGFP1 and saGFP2 constructs. Only those protoplasts that were co-transformed with C-terminal fusions of saGFP2 to KEA3 or Alb3 and stromal localized saGFP1 (transit peptide alone or cpSRP43 fusion), yielded GFP fluorescence. Scale bar, 15 µm.

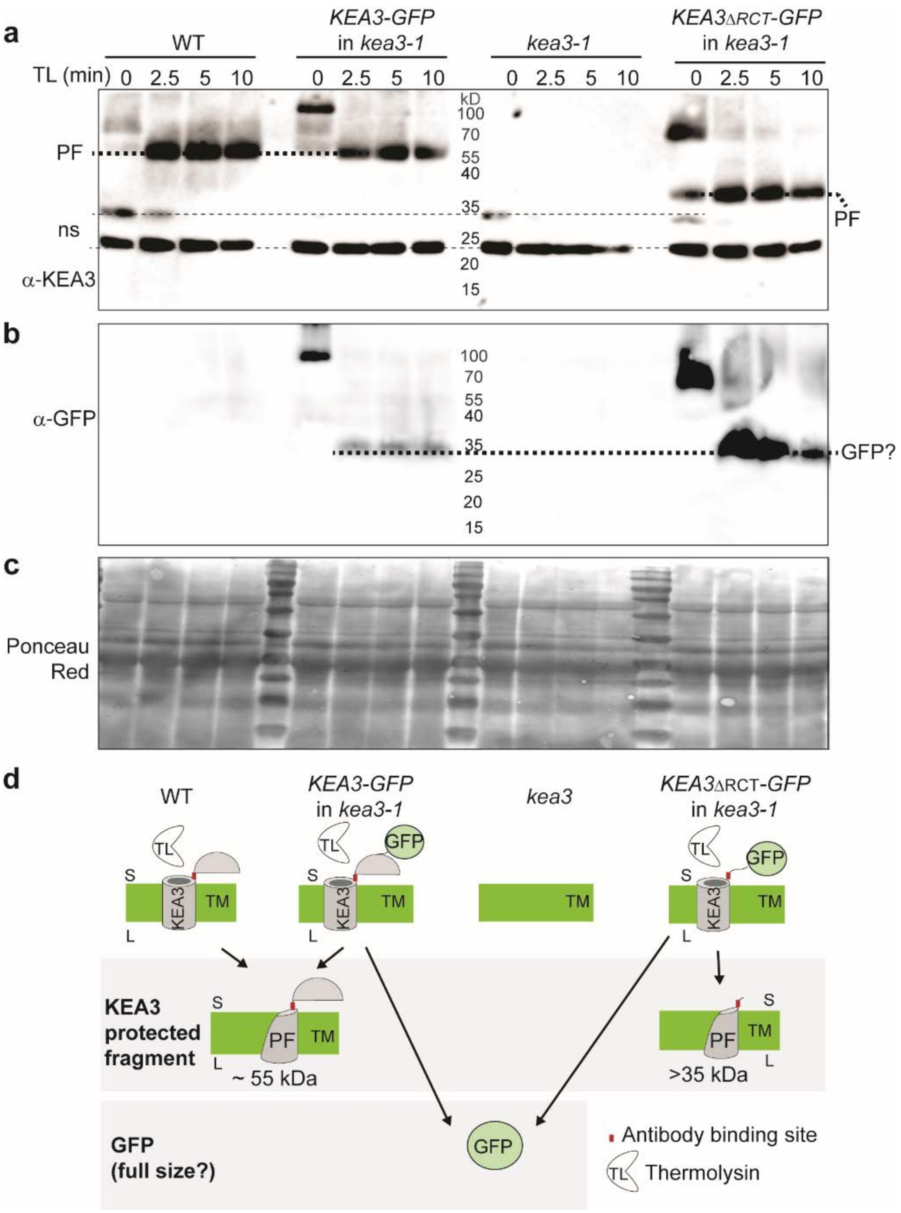


**Supplemental Fig. 2 Thermolysin digestion results of intact thylakoids can be interpreted by a stromal localization of the RCT**

**a-b**, Intact thylakoids from WT, *KEA3-GFP* in *kea3-1*, *kea3-1* and *KEA3*$_{\Delta RCT}$*-GFP* in *kea3-1* were treated with thermolysin and samples were taken at 0, 2.5, 5 and 10 min after addition of the protease. Proteins were separated by SDS-PAGE and detected by immunoblotting using a KEA3 (a) or GFP-specific (b) antibody. **c**, Ponceau Red staining of the membrane after blotting. **d**, Model of a stromal localized RCT in line with the results shown in a-b. Thylakoids harboring either the native KEA3 or KEA3-eGFP showed a protected fragment of the same molecular weight after the thermolysin treatment, in line with a stromal exposed RCT being cut off by thermolysin. Thylakoids harboring a KEA3$_{\Delta RCT}$-eGFP version showed a protected smaller fragment, suggesting that the protected fragment in KEA3-eGFP comprises at least part of the C-terminus. Detection of eGFP showed fragments of a size corresponding to full-length eGFP, for both KEA3-eGFP and KEA3$_{\Delta RCT}$-eGFP after digestion. This may be due to the release of stromal eGFP from intact thylakoids by the thermolysin treatment, which would assume eGFP to be resistant to further fragmentation by thermolysin in the reaction mix. Alternatively, the fragment detected by the GFP antibody may represent two different hybrids of similar size consisting of part of GFP and fragments of either KEA3 or KEA3$_{\Delta RCT}$, respectively. (TL, thermolysin; PF, protected fragment; TM, thylakoid membrane; S, stroma and L, lumen, ns, nonspecific signal)

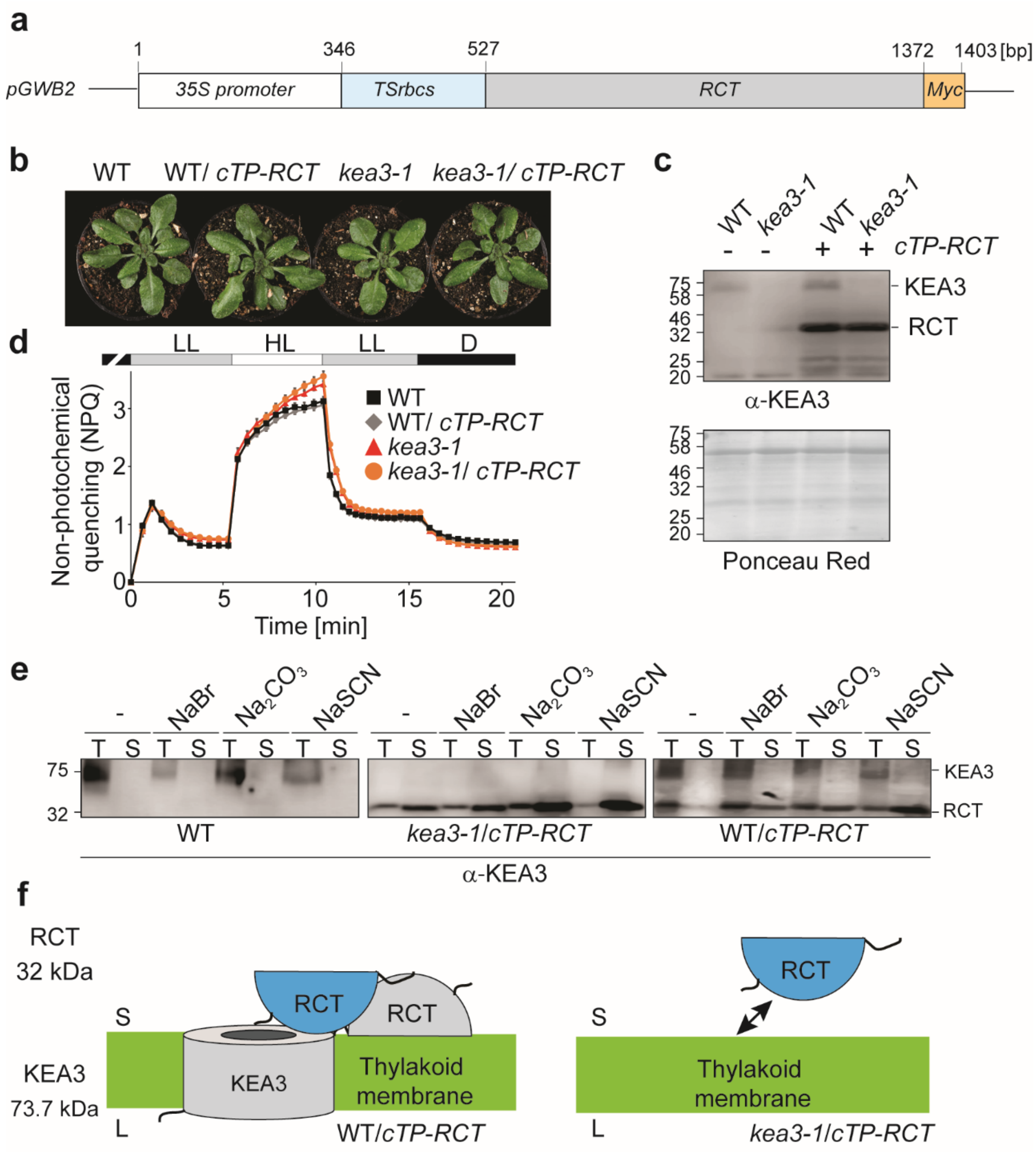


**Supplemental Fig. 3 Native KEA3 stabilizes stromal RCT**

**a**, Construct that targets the Myc-tagged KEA3 regulatory C-terminus (RCT, AAs 495-776) to the chloroplast stroma (*TSrbcs*, sequence for chloroplast targeting of Arabidopsis RbcS3B; *pGWB2*, binary vector used for cloning). **b**, Picture of four-week-old WT, *kea3-1* and *cTP-RCT* expressing plants. **c**, Immunoblot analysis of total protein extract from plants shown in b, using a KEA3 specific antibody, reveals *cTP-RCT* plants to accumulate high levels of additional RCT. The Ponceau Red stained membrane prior to detection is shown as a loading control. **d**, Non-photochemical quenching (NPQ) analysis of 30 min dark acclimated plants as in b, exposed to 5 min of low light (LL: 90 µmol photons $m^{-2}$ $s^{-1}$), 5 min high light (HL: 900 µmol photons $m^{-2}$ $s^{-1}$), 5 min LL and 5 min darkness (D). **e**, Immunoblot analysis of thylakoid (T) and soluble (S) fractions from isolated thylakoids incubated with buffer alone (-), 2 M NaBr, 0.1 M $Na_2CO_3$ or 2 M NaSCN from plants as indicated below the blotting results using a KEA3 specific antibody. **f**, Model explaining results in e. In the presence of KEA3 in the thylakoid membrane stromal RCT is stabilized via protein-protein interactions. In the absence of KEA3, RCT has some capacity to interact with the thylakoid membrane, but is rapidly washed off with buffer (RCT, regulatory C-terminus of KEA3; S, stroma and L, lumen).

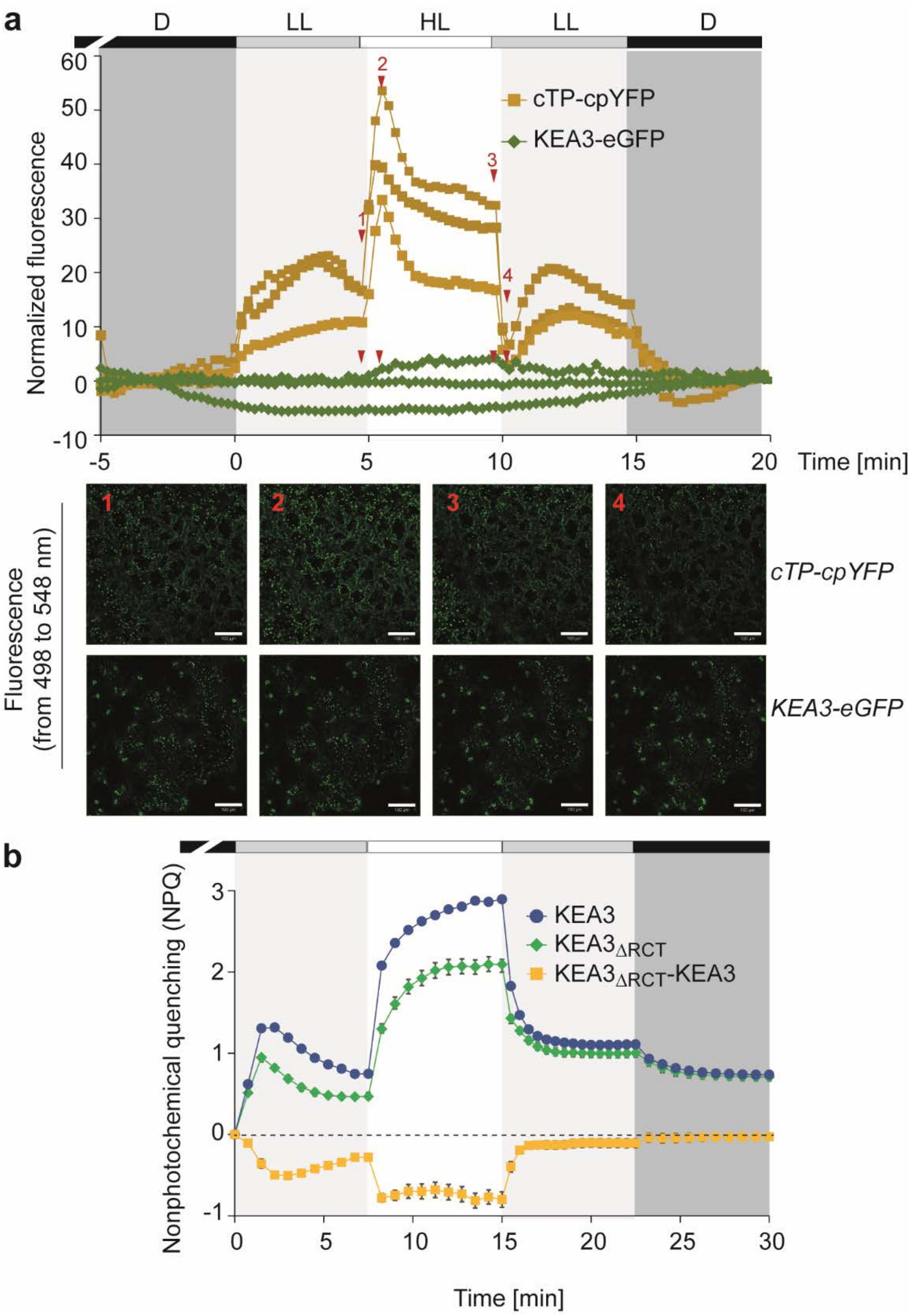


**Supplemental Fig. 4 pH transients and KEA3 regulation coincide.**

**a**, Three independent baseline-subtracted fluorescence recordings of plants harboring either a chloroplast targeted cpYFP or KEA3-eGFP used to generate the graph shown in Fig. 2a (D, darkness; LL, low light of 90 µmol photons $m^{-2}$ $s^{-1}$; HL, high light of 900 µmol photons $m^{-2}$ $s^{-1}$). Numbered arrows in red indicate time points at which representative microscopy pictures are displayed below the graph. **b**, Plants expressing WT-levels of KEA3 and $KEA3_{\Delta RCT}$ (KEA3-GFP in *kea3-1* and $KEA3_{\Delta RCT}$ -GFP in *kea3-1*, respectively) were exposed to the same fluctuating light treatment as in a and NPQ was calculated from Chl *a* fluorescence. The NPQ difference between $KEA3_{\Delta RCT}$ and KEA3 was calculated and reflects the loss of NPQ in $KEA3_{\Delta RCT}$ due to the absence of proton antiport regulation via the RCT. Average is shown for n = 5 ± SD.

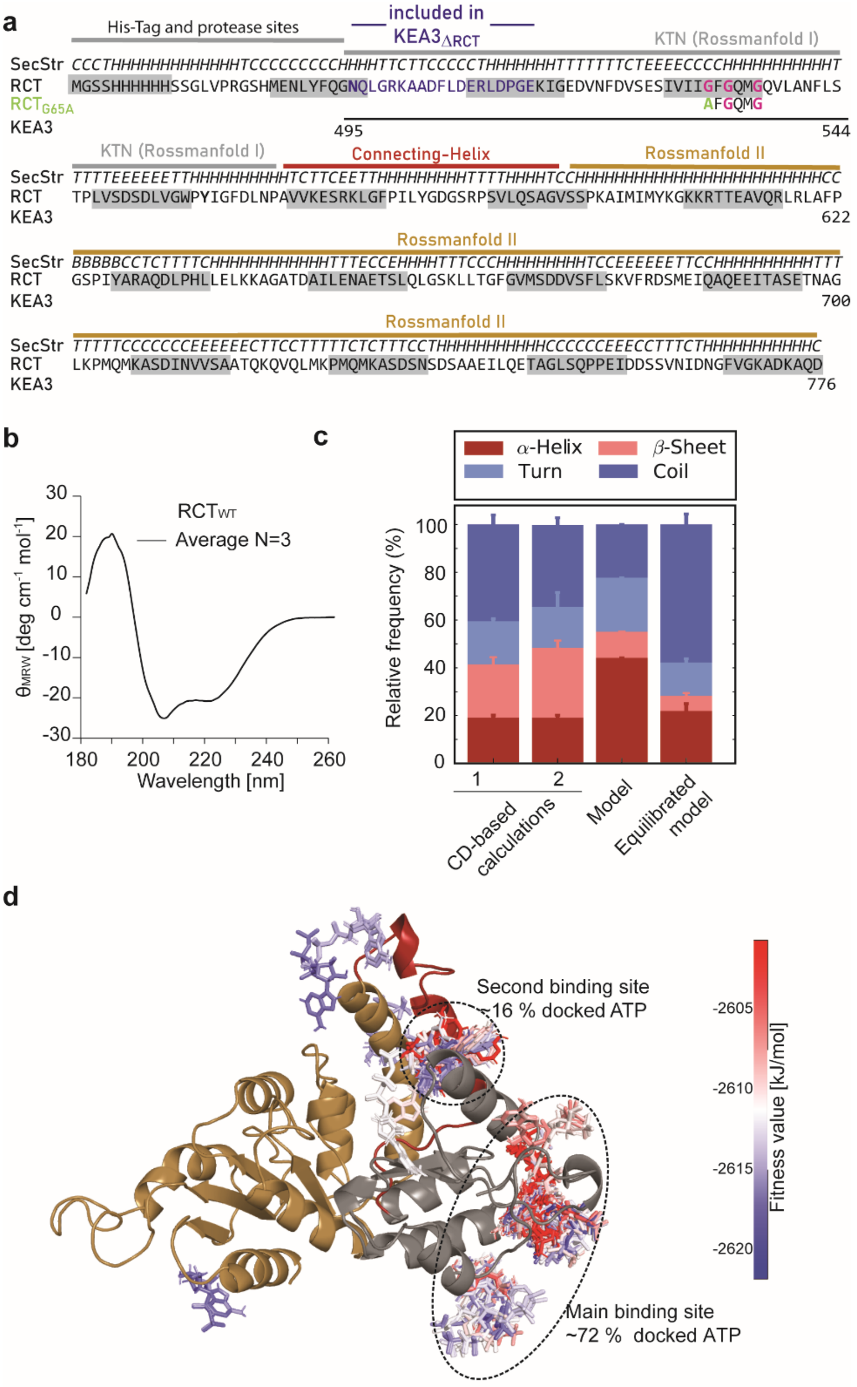


**Supplemental Fig. 5 RCT sequence and structure, experimental verification and ATP docking**

**a,** Sequence, predicted secondary structure and folds of the recombinant RCT with its N-terminal His-tag. Blue highlights the sequence retained in KEA3$_{\Delta RCT}$ and start of RCT. The conserved glycine stretch is marked in purple, the alanine replacement at G65 in green. The RCT spans KEA3 residues 495-776 (SecStr, secondary structure: C, coil; T, turn; H, α-helix; E, ß-sheet; B, ß-bridge). **b**, Far-UV spectrum of recombinant RCT. Average of n=3 is shown **c**, Secondary structure composition as estimated from the experimental data in (b) according to reference spectra of native globular proteins (1) and denatured proteins (2) with n = 3, the model built by MODELLER and the equilibrated model after 1 µs molecular dynamics simulation with n = 10 simulations. Error bars indicate SD. **d**, The RCT model was used for in silico docking experiments using SwissDock and applying 256 different ATP structures. Docking results are ranked according to SwissDock's fitness value with blue indicating the strongest interaction.

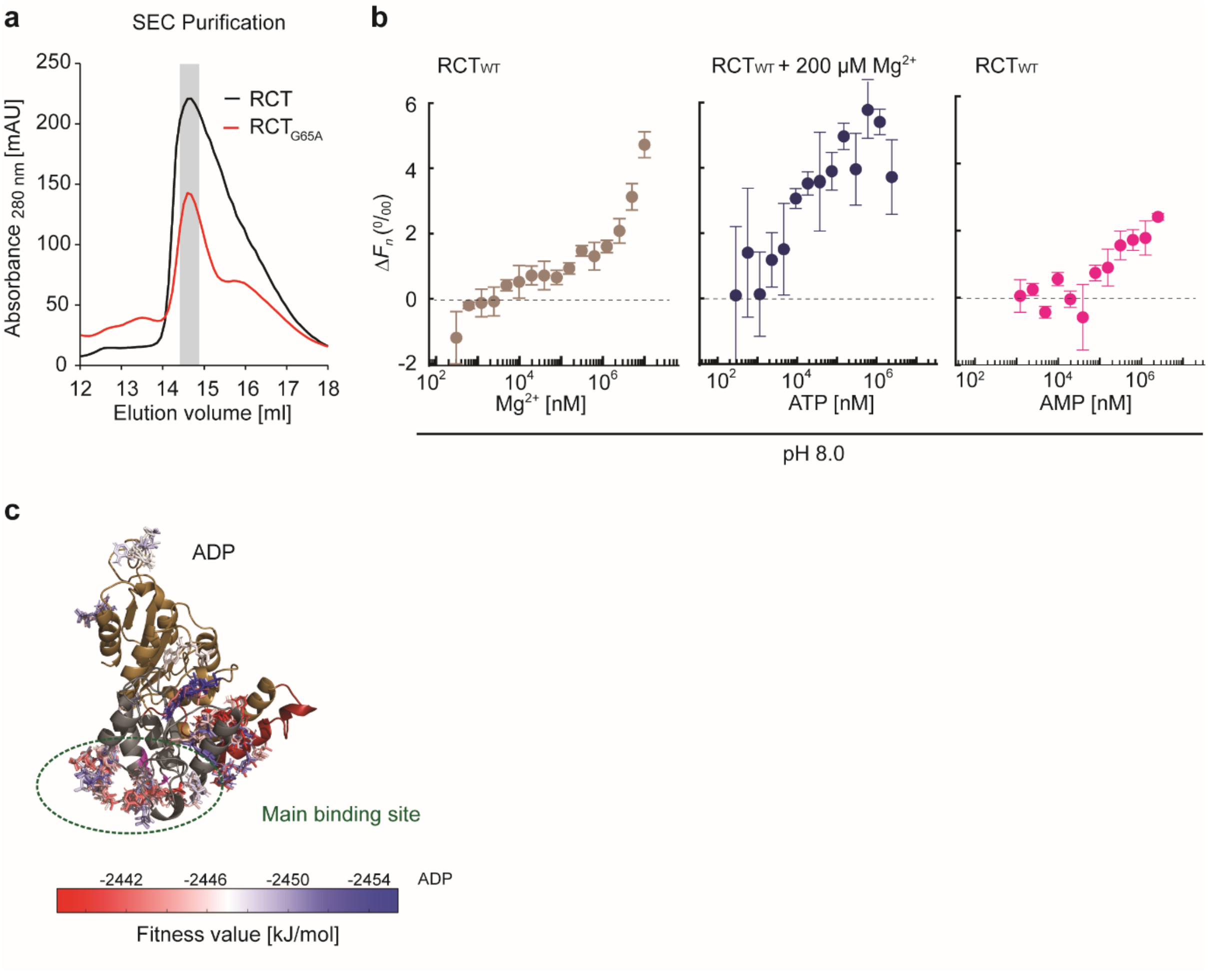


**Supplemental Fig. 6**

**a**, Recombinant RCT and $RCT_{G65A}$ were purified via size exclusion chromatography and eluted at the same elution volume of 14.5 – 15.0 ml for subsequent nucleotide binding analyses. **b**, Additional MST analyses of the $RCT_{WT}$ at pH 8.0 to test effect of $Mg^{2+}$ on ATP binding and AMP binding. **c**, *In silico* docking analysis using 256 different ADP structures places ADP into the same binding pocket as ATP. Docking results are ranked according to SwissDock's own fitness value with blue indicating the strongest interaction. 44 % of ADP molecules docked to the main binding site close to the glycine stretch.

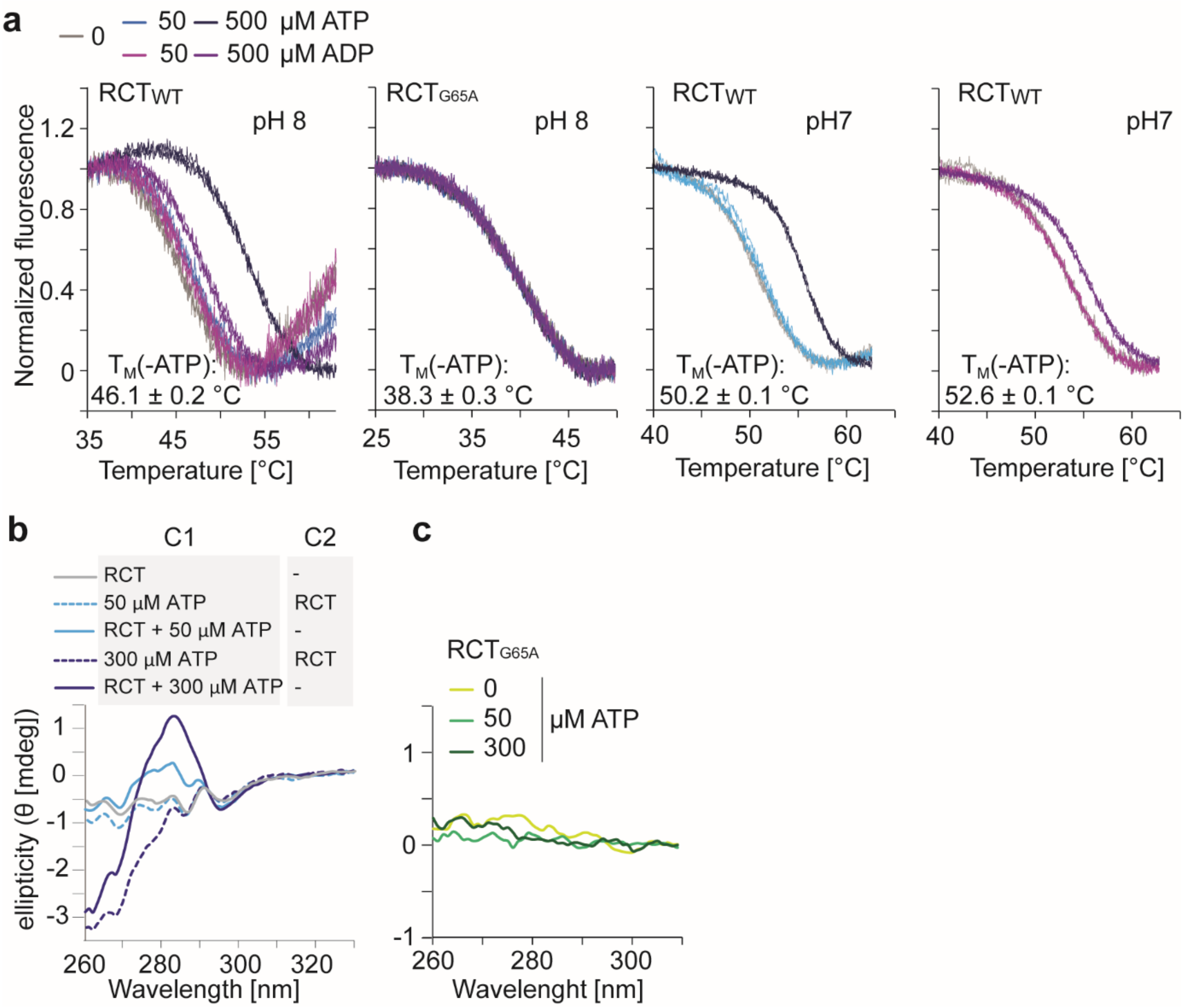


**Supplemental Fig. 7 Upon ATP binding the RCT undergoes a conformational change, which results in increased thermal stability and differences in CD spectra**

**a**, Differential scanning fluorimetry was used to record melting curves for analyzing effects of ATP or ADP binding on the thermal stability of RCT and $RCT_{G65A}$ at pH 7.0 and pH 8.0. Single traces for n = 3 are shown. The melting temperature ($T_M$) was derived from the minimum of the first derivative of the melting curve. **b**, Near-UV CD spectra were recorded with a two-cuvette set-up (C, cuvette) and revealed that RCT undergoes a conformational change in response to ATP binding at pH 8. **c**, $RCT_{G65A}$ was measured at pH 8.0 with different concentrations of ATP, which did not majorly effect ellipticity.

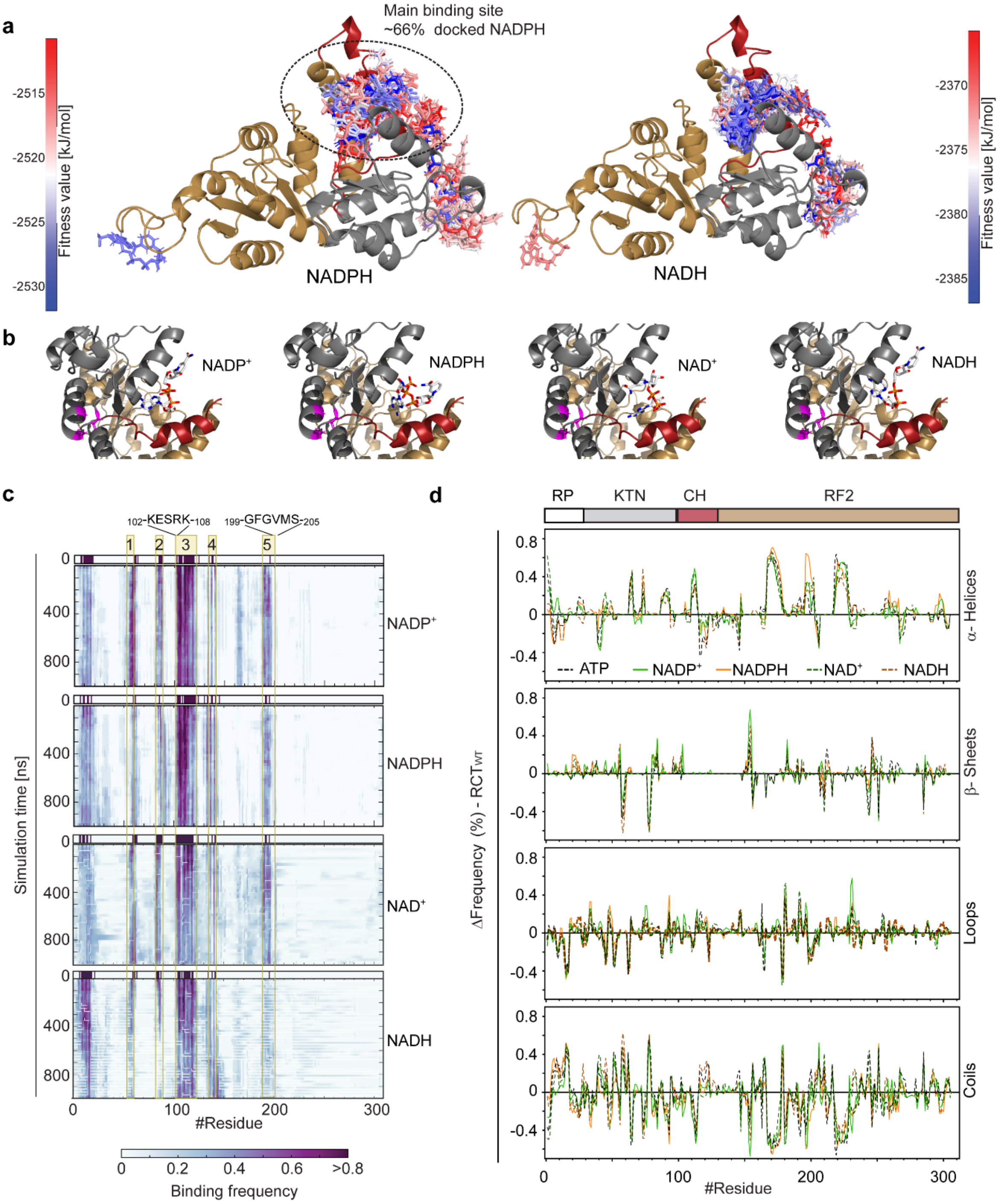


**Supplemental Fig. 8 NAD species bind to a different binding site than ATP and ADP, but have similar effects on secondary structure during simulation**

**a**, *In silico* docking analysis using 256 different structures of all four NAD-species placed these nucleotides into a different binding pocket than ATP or ADP. Docking results are ranked according to SwissDock's fitness value with blue indicating the strongest interaction and are shown for NADPH and NADH. **b**, The best fit binding site is the same for all four species. **c**, Molecular dynamics (MD) simulations were performed on $RCT_{WT}$ with NAD-species docked to their best-fit binding site. The distance from the nucleotide was calculated for each amino acid residue during the time course of the simulation. Binding frequency was defined by a distance of less than 7 Å from the nucleotide. Five nucleotide adjacent regions were identified through the docking with region 3 containing a stretch of positively charged amino acids and region 5 containing a modified glycine-rich stretch. **d**, Simulated differences in secondary structure frequencies between free and nucleotide-docked versions of RCT. Frequencies are calculated from n = 10 simulations.

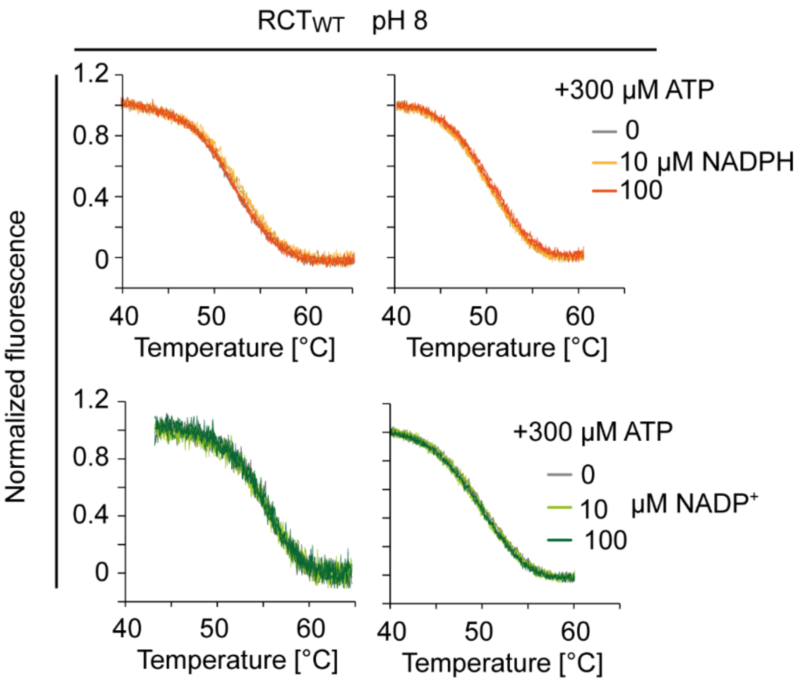


**Supplemental Fig. 9 NADPH and NADP$^{+}$ have no effect on thermal stability of the RCT$_{WT}$**

Differential scanning fluorimetry was used to record melting curves for analyzing effects of NADPH or NADP+ on the thermal stability of RCT at pH 8.0 with and without saturating concentrations of ATP. Single traces for n = 3 are shown. Neither NADPH nor NADP$^{+}$ appeared to influence thermostability under the analyzed conditions.

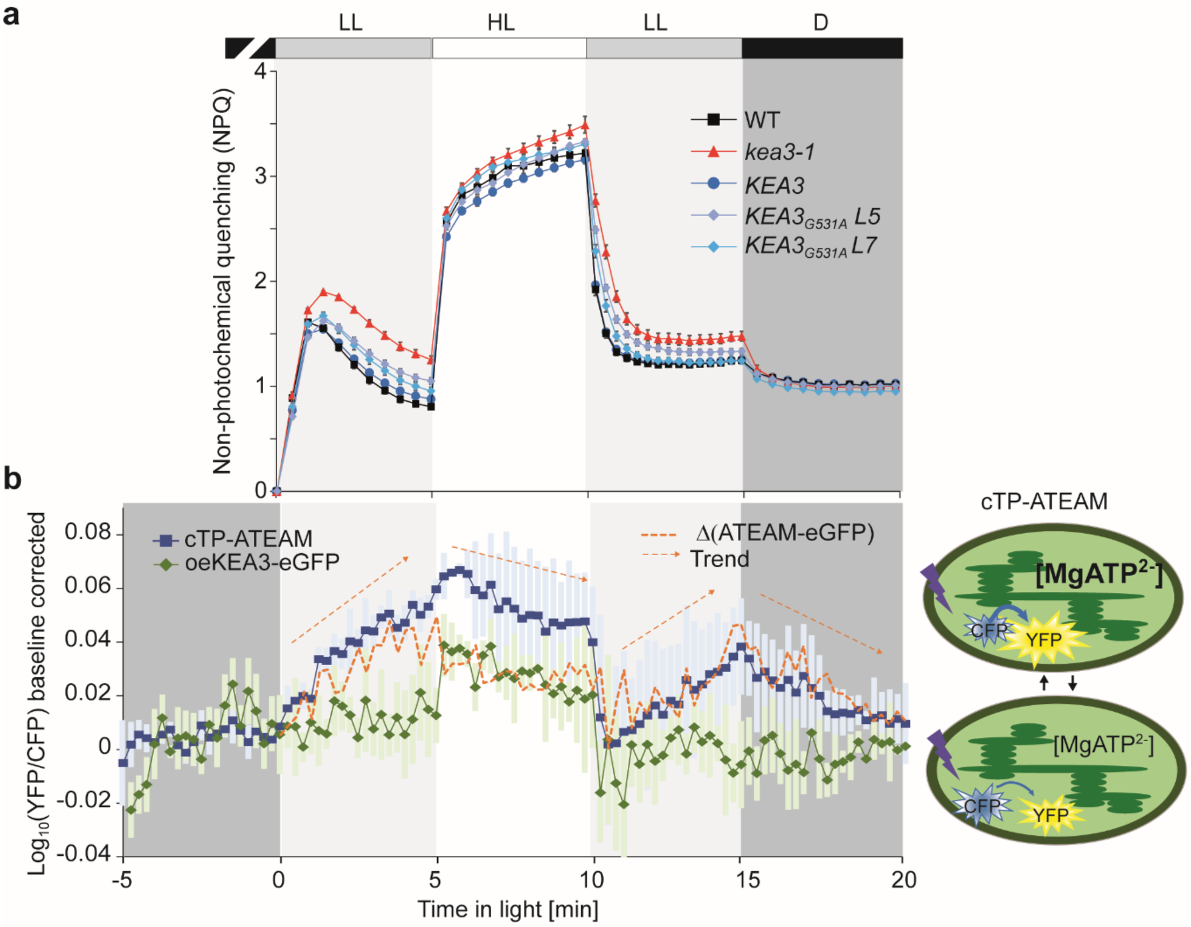


**Supplemental Fig. 10 NPQ of different lines and $MgATP^{2-}$ measurements during changes in light intensity**

**a,** The different lines were dark acclimated for 30 min dark acclimation and then exposed to an alternating light regime of 5 min low light (LL, 90 µmol photons $m^{-2}$ $s^{-1}$), 5 min high light (HL, 900 µmol photons $m^{-2}$ $s^{-1}$), 5 min LL and 5 min darkness and non-photochemical quenching (NPQ) was determined. **b**, Leaf discs of cTP-ATeam or oeKEA3-eGFP (control) were dark-acclimated (D, darkness) and examined by confocal microscopy using a light box, which supplied red light between 620 and 645 nm wavelengths at two different light intensities (low light, LL, 90, and high light, HL, 900 µmol photons $m^{-2}$ $s^{-1}$) during indicated time intervals. Excitation was provided at 458 nm and fluorescence was collected for CFP at 470 – 507 nm and for YFP at 521 – 531 nm. The fluorescence signals for YFP/CFP were log10- transformed and a baseline from the last minute of each dark phase (before and after the light treatment) was subtracted. Average is shown for n = 3 ± SD. Note that also the YFP/CFP ratio of the control shows some reaction to changes in light intensity. The dashed line shows the average normalized ATeam signal subtracted from the normalized average control signal. Based on the behavior of the differences, arrows were placed to indicate the trend of the normalized ATeam signal.

**Supplemental Table 1** Vectors generated for self-assembling GFP analysis

Coding sequences of *KEA3*, *cpSRP43*, ALB3 or the transit sequence of the small rubisco subunit (TSrbcs) were amplified from cDNA and inserted into *Kpn*I and *Spe*I restriction sites of pAVA-saGFP1-10, pAVA-saGFP11N or pAVA-saGFP11C resulting in saGFP1-10 (large GFP fragment; saGFP1) or N- or C-terminal saGFP11 (small GFP fragment; saGFP2) fusions, respectively[14].
The plasmid pAVA-saGFP11N was further modified introducing two additional restriction sites (*Bgl*II and *Xma*JI) upstream of saGFP11 using site directed mutagenesis (SDM). These restriction sites were used to introduce the transit peptide of the RubisCo small subunit (TSrbcs; *At5g38410*) leading to the plasmid pAVA-TSrbcs-saGFP11N. Mature KEA3 was introduced into this plasmid downstream of saGFP11 resulting in a TSrbcs-saGFP11-KEA3 fusion (saGFP2 N-KEA). Additionally, precursor KEA3 containing the chloroplast targeting peptide was cloned into pAVA-saGFP11C resulting in a C-terminally fused saGFP11 (saGFP2 KEA-C). As a control, precursor *ALB3* cDNA was cloned into pAVA-saGFP11C (saGFP2 Alb3-C). The chloroplast localization of the large saGFP1-10 fragment was ensured either by C-terminal fusion to TSrbcs (saGFP1 Str) or to TSrbcs-cpSRP43 (saGFP1 Str-SRP43-C). TSrbcs was inserted via in-fusion cloning according to manufacturer's instructions (Takara) into the *Kpn*I linearized pAVA-saGFP1-10 plasmid. TSrbcs-cpSRP43 was introduced into *Kpn*I and *Spe*I restriction sites of pAVA-saGFP1-10.

| # | Plasmid | Information | Primer sequences used (5' → 3') |
|---|---|---|---|
| 1 | pAVA-saGFP11N* | To obtain N-terminal saGFP2 fusions | |
| 2 | pAVA-saGFP11N BglII | *Bgl*II site introduction into #1 via SDM | GCAGCAATTTAAATCAGATCTTTTAAAGCAAAAGC<br>GCTTTTGCTTTAAAAGATCTGATTTAAATTGCTGC |
| 3 | pAVA-saGFP11N BglII, XmaJI | *Xma*JI site introduction into #2 via SDM | GCAATTTTCTGAAAATTTTCACCTAGGACGAACGATA GCCATGCG<br>CGCATGGCTATCGTTCGTCCTAGGTGAAAATTTTCA GAAAATTGC |
| 4 | pAVA-TSrbcs-saGFP11N | TSrbcs introduction into #3 | TAGAGATCTATGGCTTCTATGATA<br>CTACCTAGGTACTTTACTCTTCCACC |
| 5 | pAVA-TSrbcs-saGFP11-KEA3 | Introduction of mature *KEA3* into #4 to obtain saGFP2-N-KEA3 | s. Supplemental Table 3 |
| 6 | pAVA-saGFP11C* | To obtain C-terminal saGFP2 fusions | |
| 7 | pAVA-KEA3-saGFP11 | Introduction of full length *KEA3* into #6 to obtain KEA3-C-saGFP2 | s. Supplemental Table 3 |
| 8 | pAVA-Alb3-saGFP11 | Introduction of full length *Alb3* into #6 | GCTGGTACCATGGCGAGAGTTCTA<br>AGCACTAGTTACAGTGCGTTTCCG |
| 9 | pAVA-saGFP1-10* | For cytosolic saGFP1 | |
| 10 | pAVA-TSrbcs-saGFP1-10 | For stromal saGFP1 | GCAGCAATTTAAATCAGATCTATGGCTTCTATGATA<br>TAGTCATGCGGCCGCGGTACCACTTTACTCTTCCAC |
| 11 | pAVA-TSrbcs-cpSRP43-saGFP1-10 | Introduction of full length *cpSRP43* into #10 to obtain cpSRP43-C-saGFP1 | TAGGGTACCATGGCTTCTATGATA<br>CTAACTAGTTTCATTCATTGGTTGTTGTTG |

* kindly provided by E. Schleiff (Frankfurt, Germany), containing sequences provided by G. S. Waldo (Los Alamos, NM, USA)[15]

**Supplemental Table 2**: Further primer sequences used in this study.

| **Primer sequences (5' → 3')** | **Purpose** |
|---|---|
| TGGCACAGGTTCTGGCCAATTTTTTGTCAACGC | Site-directed mutagenesis to derive $RCT_{G65A}$ |
| TTTGTGCAAATGCAATGATGACAATTGATTCACTAAC | |
| TTGGCATGGGACTAACTCAG | Genotyping of $KEA3_{G531A}$ plants |
| CACAATCAGACCACCCAAGA | |
| TCATTTGGAGAGAACACGGGGGACTCTAGATGGCTTC TATGATATCCTCTTC | Amplification of TSrbcs (*At5g38410*) for stromal RCT overexpression using the *pGWB2* vector |
| TTCCAAGTTGGTTCTTTACTCTTCCACCATTGC | |
| TGGAAGAGTAAAGAACCAACTTGGAAGAAAAG | Amplification of RCT with C-terminal Myc-tag for overexpression using the *pGWB2* vector |
| TTGAACGATCGGGGAAATTCGAGCTCTTACAGATCCTC TTCTGAGATGAGTTTTTGTTCATCTTGAGCTTTATCAGC TTTAC | |
| ATCAAGCATTCTACGGGTACCATGGCAATTAGTACTAT GTTAG | Amplification of the full length *KEA3* coding sequence without stop codon for insertion upstream of the saGFP2 coding sequence in the pAVA vector for transient expression of KEA3-C-saGFP2. |
| CAGACCCTCCATCGGATCCACTAGTTGATATCACCACT TTGTAC | |
| CAATCACCGGATTGGGGTACCATGGCAATTAGTACTAT GTTAG | Amplification of the *KEA3* coding sequence with stop codon for insertion downstream of TSrbcs and the *saGFP2* coding sequence in the pAVA vector for transient expression of TSrbcs-saGFP2-N-KEA3. |
| GATTTTTGCGGACTCTAGATTAACTAGTTTAATCTTGAG CTTTATCAGC | |
| TTGGCATGGGACTAACTCAG | Genotyping of the *kea3-1* mutant (Gabi_170G09) |
| CACAATCAGACCACCCAAGA | |

### References


1 Uflewski, M. *et al.* Functional characterization of proton antiport regulation in the thylakoid membrane. *Plant Physiol*, doi:10.1093/plphys/kiab135 (2021).

2 Armbruster, U. *et al.* Regulation and Levels of the Thylakoid K+/H+ Antiporter KEA3 Shape the Dynamic Response of Photosynthesis in Fluctuating Light. *Plant Cell Physiol* **57**, 1557-1567, doi:10.1093/pcp/pcw085 (2016).

3 Behera, S. *et al.* Cellular Ca2+ Signals Generate Defined pH Signatures in Plants. *The Plant Cell* **30**, 2704-2719, doi:10.1105/tpc.18.00655 %J The Plant Cell (2018).

4 De Col, V. *et al.* ATP sensing in living plant cells reveals tissue gradients and stress dynamics of energy physiology. *eLife* **6**, doi:10.7554/eLife.26770 (2017).

5 Armbruster, U. *et al.* Ion antiport accelerates photosynthetic acclimation in fluctuating light environments. *Nat Commun* **5**, 5439, doi:10.1038/ncomms6439 (2014).

6 Nakagawa, T. *et al.* Development of series of gateway binary vectors, pGWBs, for realizing efficient construction of fusion genes for plant transformation. *Journal of bioscience and bioengineering* **104**, 34-41, doi:10.1263/jbb.104.34 (2007).

7 Clough, S. J. & Bent, A. F. Floral dip: A simplified method for Agrobacterium-mediated transformation of Arabidopsis thaliana. *Plant Journal* **16**, 735–743, doi:10.1046/j.1365-313X.1998.00343.x (1998).

8 Jorgensen, W. L., Chandrasekhar, J., Madura, J. D., Impey, R. & Klein, M. L. J. J. o. C. P. Comparison of simple potential functions for simulating liquid water. **79**, 926-935 (1983).
9 Bussi, G., Donadio, D. & Parrinello, M. Canonical sampling through velocity rescaling. **126**, 014101, doi:10.1063/1.2408420 (2007).
10 Parrinello, M. & Rahman, A. Polymorphic transitions in single crystals: A new molecular dynamics method. **52**, 7182-7190, doi:10.1063/1.328693 (1981).
11 Hess, B., Kutzner, C., van der Spoel, D. & Lindahl, E. GROMACS 4: Algorithms for Highly Efficient, Load-Balanced, and Scalable Molecular Simulation. *Journal of chemical theory and computation* **4**, 435-447, doi:10.1021/ct700301q (2008).
12 Essmann, U. *et al.* A smooth particle mesh Ewald method. *The Journal of Chemical Physics* **103**, 8577-8593, doi:10.1063/1.470117 (1995).
13 Sreerama, N. & Woody, R. W. Estimation of protein secondary structure from circular dichroism spectra: comparison of CONTIN, SELCON, and CDSSTR methods with an expanded reference set. *Anal Biochem* **287**, 252-260, doi:10.1006/abio.2000.4880 (2000).
14 Ulrich, T., Gross, L. E., Sommer, M. S., Schleiff, E. & Rapaport, D. Chloroplast β-barrel proteins are assembled into the mitochondrial outer membrane in a process that depends on the TOM and TOB complexes. *J Biol Chem* **287**, 27467-27479, doi:10.1074/jbc.M112.382093 (2012).
15 Cabantous, S., Terwilliger, T. C. & Waldo, G. S. Protein tagging and detection with engineered self-assembling fragments of green fluorescent protein. *Nature Biotechnology* **23**, 102-107, doi:10.1038/nbt1044 (2005).
